# Tet1 safeguards lineage allocation in intestinal stem cells

**DOI:** 10.1101/2025.05.08.652522

**Authors:** Antoine Gleizes, Nicolas V Janto, Sergei Bombin, Vivek Rao, Gurel Ari, Siyang Sun, Maria Fonseca, Adam D Gracz

## Abstract

Intestinal stem cells (ISCs) balance self-renewal and differentiation to maintain the intestinal epithelial barrier, which is replaced weekly throughout adult life. Genetic control of ISC differentiation is well-defined relative to transcription factor (TF) activity, but less is known regarding the role of chromatin regulation in ISC biology. Prior work from our lab and others has shown that *Tet1*, a chromatin modifying enzyme involved in DNA demethylation, is specifically enriched in ISCs and early secretory progenitors. While constitutive loss of *Tet1* is associated with defects in early postnatal ISC development, its role in adult ISC biology remains unknown. Here, we show that *Tet1* safeguards ISC fate decisions by reducing sensitivity to extrinsic signaling. Inducible, intestine-specific *Tet1* knockout mice (Tet1iKO) exhibit environmentally sensitive phenotypes, including absorptive differentiation bias and premature expression of mature absorptive transcripts in ISCs. These phenotypes are largely “silenced” in animals housed in a high-level barrier facility, where Tet1iKO epithelium closely resembles controls. Despite the lack of baseline phenotype in these conditions, Tet1iKO mice retain increased sensitivity to pro-differentiation signaling. *In vivo*, succinate administration induces increased tuft and goblet cell hyperplasia in the absence of *Tet1*, while Tet1iKO organoids cultured with IL-4 or DAPT exhibit increased tuft and enteroendocrine cell specification, respectively. While ATAC-seq of Tet1iKO ISCs reveals minimal changes in chromatin accessibility, footprinting analysis suggests increased binding of lineage-specific TFs and CTCF even in the absence of cellular phenotypes. Together, our data demonstrate that *Tet1* serves as a “buffer” against ISC differentiation and suggest that it does so in a lineage agnostic manner that is not dependent on changes in chromatin accessibility.

## Introduction

Intestinal stem cells (ISCs) reside at the base of crypt glands and divide approximately once every 12-24hrs to give rise to transit-amplifying progenitors (TAs), which undergo further differentiation to replace mature intestinal epithelial cells (IECs). This continual renewal results in near-total turnover of mature IECs every 5-7 days. Cell fate specification of ISCs must be tightly regulated to ensure the appropriate balance of mature absorptive and secretory cell numbers, which can change in response to extrinsic environmental factors like inflammation, injury, or infection. ISC differentiation is well characterized relative to a host of lineage-specific transcription factors (TFs) activated in response to upstream signaling. For example, secretory vs. absorptive fate represents the earliest step in terminal differentiation of ISCs and is regulated by the opposing activity of TFs ATOH1 and HES1 ^1,2^. HES1 is induced by Notch signaling and in turn represses ATOH1, while Notch inhibition downregulates HES1 to derepress ATOH1 ^1^. More broadly, WNT signaling generally promotes stemness while BMP signaling promotes differentiation. Further specification of mature IECs, including absorptive, goblet, Paneth, enteroendocrine, and tuft cell populations, has been extensively studied through hierarchical activation of cell type specific signaling pathways and their effector TFs (reviewed in ^3^).

While our understanding of *trans*-regulatory networks controlling ISC fate has grown with improved biomarkers of IEC subpopulations and single cell transcriptomics, the role of the *cis*-regulatory chromatin landscape has proven harder to define. Early studies of chromatin in the intestine proposed a model where ISCs and their progeny have highly similar or “broadly permissive” chromatin landscapes, with cell fate regulation depending largely on TFs ^4^. Evidence for this “*trans*”-dominant model of regulation has been shown for closely related Paneth and goblet cell lineages, which share accessible chromatin landscapes and appear to be specified entirely by TFs in response to differential BMP and WNT signaling ^5^. Transient ISC ablation also results in a damage response that involves little discernible remodeling of chromatin in other cell types ^6^. However, other studies have demonstrated significant changes in chromatin accessibility with ISC differentiation, especially for secretory cells ^6–9^. Further, irradiation injury induces facultative stem cell function in secretory progenitors, which undergo substantial chromatin remodeling as they reacquire an ISC state ^7^.

Our lab and others have shown that the chromatin modifying enzyme *Tet1* is specifically enriched in *Lgr5*+ ISCs ^8,10^. *Tet1* belongs to the Ten-eleven translocation (TET) family, consisting also of *Tet2* and *Tet3*, which are expressed broadly across all IECs without cell type specific enrichment ^8^. TETs play diverse and context-dependent roles in chromatin regulation and were first characterized by their ability to convert 5-methylcytosine (5mC) to 5-hydroxymethylcytosine (5hmC), initiating an iterative oxidation process leading to active DNA demethylation and derepression of target genes ^11,12^. Additionally, all TETs can co-recruit other regulatory complexes, with TET1 notably recruiting repressive HDAC and PRC2 complexes to silence gene expression independent of its catalytic function ^13,14^. TET1 is required for embryonic stem cell potency and facilitates reprogramming of adult cells to an induced pluripotent state, linking its broad regulatory impact to stem cell function ^11,15^.

Studies using *Tet1*-null mice have previously shown that constitutive deletion of *Tet1* results in decreased WNT signaling and reduced numbers of *Lgr5*+ ISCs in postnatal intestines ^10^. However, the role of *Tet1* in adult ISCs and whether these effects are epithelial autonomous remain unknown. Here, we generated IEC-specific, inducible *Tet1* knockout mice (Tet1iKO) to determine the regulatory impact of *Tet1* on ISC differentiation. By assessing ISC differentiation *in vivo* and *in vitro*, we find that Tet1iKO phenotypes are sensitive to extrinsic signaling, including housing conditions, which can induce or silence differentiation defects in Tet1iKO mice. Our data support a model where TET1 safeguards ISC fate decisions by “buffering” their response to changes in the extrinsic signaling environment.

## Results

### Environmental conditions influence intestinal epithelial-specific Tet1 knockout phenotype

To study the role of *Tet1* in adult ISCs, we generated an inducible, intestinal epithelial-specific knockout mouse model (Tet1^fl/fl^:vil^CreER^), referred to here as “Tet1iKO” ^16,17^. We induced recombination in both control (Tet1^+/+^:vil^CreER^) and Tet1iKO mice 8-24wks of age by administering tamoxifen once daily for three days, followed by two days of washout (Figure S1A). Loss of *Tet1* was validated by RT-qPCR from FACS-isolated ISCs (Figure S1B). Because *Tet1* has previously been implicated in lineage allocation of embryonic stem cells, we first assessed Tet1iKO intestines by immunofluorescence (IF) on histological sections of jejunal tissue ^11,19^. Conventionally housed Tet1iKO animals [CONV (UNC)] exhibited a significant decrease in villus-localized CHGA+ enteroendocrine cells (Figure S2A) and DCLK1+ tuft cells in both the crypts and villi (Figure S2B). Villus-localized MUC2+ goblet cells were significantly increased (Figure S2C), while numbers of LYZ+ Paneth cells remained unchanged (Figure S2D). Because mice carrying constitutive null alleles for *Tet1* were reported to have decreased ISC numbers, we quantified OLFM4+ cells and noted significant increase in Tet1iKO intestines (Figure S2E). Together, these data demonstrate that loss of *Tet1* in adult intestinal epithelium results in subtle but significant changes in multiple secretory cell types and may increase ISC numbers.

Over the course of these studies, our lab moved to a new institution. Because our mice were housed in a new facility [CONV (Emory)], we repeated histological quantification to validate the Tet1iKO phenotype (Figure 1A-D and S2F-G). While goblet cell numbers remained significantly increased in the new facility (Figure 1A & E), we observed substantial changes in our previously identified Tet1iKO phenotype. Enteroendocrine (Figure 1B & F) and tuft cell numbers (Figure 1C & G), previously significantly reduced, were unchanged but trending upward in Tet1iKO mice housed in the new facility. Paneth cells were also unchanged between Tet1iKO and control crypts, consistent with our former facility (Figure S2F). However, the increase in OLFM4+ cells observed in CONV (UNC) housed mice was not found in the new facility (Figure S2G). Additionally, we noted a significant enrichment for FABP1 staining further down the villus axis, suggesting increased enterocyte differentiation or changes in proportions of enterocyte populations in Tet1iKO intestines (Figure 1D & H). Interestingly, tuft cell numbers in both control and Tet1iKO mice were elevated ∼2-3 fold in our new facility, suggesting an overall change in secretory specification associated with environmental conditions. Prior reports have shown that the common commensal protist, *Tritrichomonas muris* (*Tmu*), is capable of inducing tuft cell hyperplasia through a type 2 immune response ^20–24^. Observation of cecal contents by light microscopy confirmed the presence of *Tmu* in the digestive tract of control and Tet1iKO mice in our new facility (Figure S2H). Taken with our initial phenotyping, data from CONV housed mice suggest that previously identified Tet1iKO phenotypes were masked by Tmu colonization.

**Figure 1.**
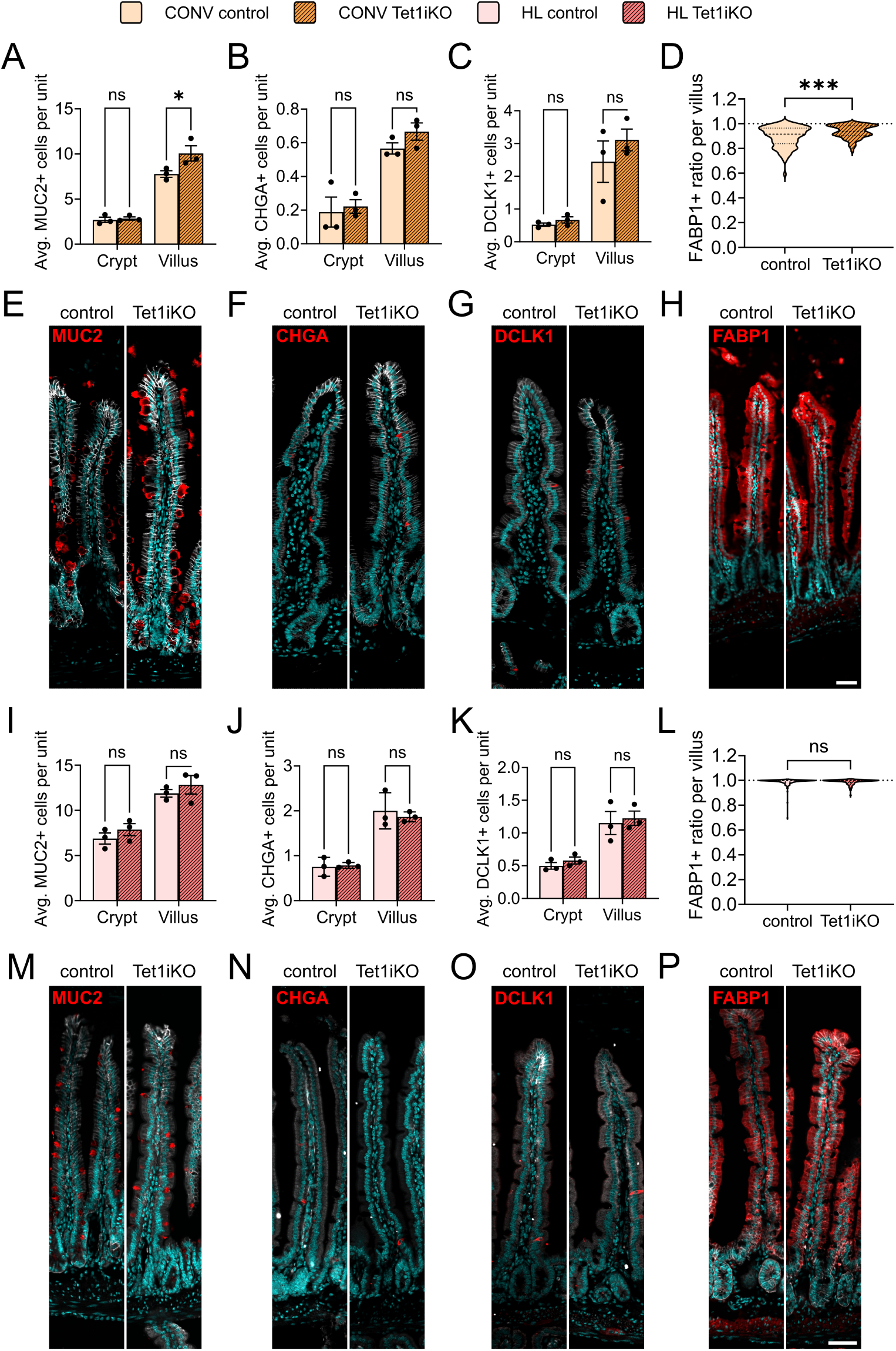
Tet1iKO promotes goblet cell and absorptive enterocyte differentiation in conventional housing. IF quantification on histological sections comparing CONV Tet1iKO to control jejunum reveals (A, E) a significant increase in goblet cell (MUC2) number only in villi, (B, F) no significant difference in enteroendocrine cell (CHGA) number, and (C, G) no significant difference in tuft cell (DCLK1) number. (D, H) Mature absorptive enterocyte marker, FABP1, exhibits a significant increase in signal length down the villus axis for Tet1iKO mice in CONV housing (a ratio of 1 equals to a positive FABP1 signal covering the full villus length, values are plotted as individual average between left and right side ratio for each villus). HL housed Tet1iKO mice exhibit no discernible phenotypes by IF, including no significant difference in (I, M) goblet cell, (J, N) enteroendocrine, or (K, O) tuft cell number. (L, P) Ratios of FABP1+ enterocytes to villus length are unchanged between HL Tet1iKO and controls (scale bars represent 50 µm) (n = 3 control and 3 Tet1iKO mice per group; n = 50 crypt or villus units per mouse; * indicates p<0.05 and *** for p<0.001).

To reduce environmental bias and eradicate *Tmu* from our experimental conditions, we rederived our mouse colony into a high-level barrier facility [HL (Emory); from here on “CONV” refers to CONV (Emory) housed mice]. Importantly, *Tmu* is specifically excluded from the HL facility. We confirmed absence of *Tmu* colonization in rederived control and Tet1iKO mice by examining cecal contents (Figure S2I). Surprisingly, histological quantification revealed a complete absence of Tet1iKO phenotype in HL conditions (Figure 1I-L & S2J-K). Tuft cell numbers were decreased in both control and Tet1iKO mice in HL relative to CONV, but we did not observe a Tet1iKO-associated increase in goblet cell differentiation (Figure 1I & M) or decrease in enteroendocrine (Figure 1J & N) or tuft cell numbers (Figure 1K & O). Further, FABP1 expression extended down the villus axis and was non-significant between Tet1iKO and controls, suggesting that decreased FABP1-positivity in the villi of CONV controls was driven by *Tmu* colonization (Figure 1L & P). Collectively, phenotyping data from three different facilities and two institutions suggest that the impact of *Tet1* on intestinal epithelial biology is highly sensitive to extrinsic cellular environments.

### Tet1 knockout is transcriptomically silent in a controlled environment

We reasoned that the lack of discernible Tet1iKO phenotype in the HL facility presented an opportunity to understand the role of *Tet1* in ISC genetic regulation with minimal environmental influences. To compare differences associated with Tet1iKO IECs in different facilities, we conducted scRNA-seq in both CONV and HL housing conditions (n=2 control and n=2 Tet1iKO per facility). Single IECs from whole intestine (duodenum to ileum) were dissociated and isolated by FACS, then subjected to split-pool construction of single cell cDNA libraries. Following quality control and filtering to remove low quality cells and multimers, data were analyzed by standard clustering and dimensionality reduction in Seurat ^25^. To focus our analysis on *Tet1*-driven changes, each facility (CONV and HL) was analyzed separately. Data sets were assessed for equal representation of individual sample (mice) and group (control vs. Tet1iKO) distribution via UMAP (Figure S3A & B and S4A & B). No obvious sampling bias was observed in either dataset, with each sample and group evenly represented. For each dataset (CONV and HL), a 90% cut-off was arbitrary set for the cumulative variance brought by each principal component identified through principal component analysis (PCA). Following this approach, 40 PCs were retained in conventional housing and 39 in HL housing (Figure S3C and S4C). Based on 90% of the total dataset variation, we identified clusters via k-nearest neighbors’ method ^26^. Optimal cluster resolution was determined semi-empirically based on: (1) cluster stability as visualized using the Clustree package and (2) expected numbers of IEC subpopulations based on known biology ^27^. We selected a 0.3 resolution corresponding to 11 clusters in CONV housing (Figure S3D & E). In HL housing, a 0.3 resolution produced 10 clusters (Figure S4D & E). The specific IEC identity of each cluster was determined by the positive enrichment for intestinal epithelial gene signatures from previously published scRNA-seq, along with canonical biomarkers (e.g.: *Lgr5* in ISCs; *Muc2* in goblet cells) (Figure S3F-L and S4F-L, Table S1) ^28^. Identified clusters were annotated accordingly. If multiple clusters were identified as sharing similar identities, they were combined to avoid overinterpretation of cellular heterogeneity (Figure 2A & E). This analytical approach resulted in 8 high-confidence, annotated clusters in samples from each facility (Figure 2A & E), with distinct cellular identities based on IEC marker distribution across clusters (Figure 2B & F). Importantly, this approach identified all major IEC lineages: ISCs, TAs, absorptive progenitors (Abs. pro.), absorptive enterocytes (Ent.), goblet cells (GC), enteroendocrine cells (EE), Paneth cells (PC), and tuft cells (TC).

**Figure 2.**
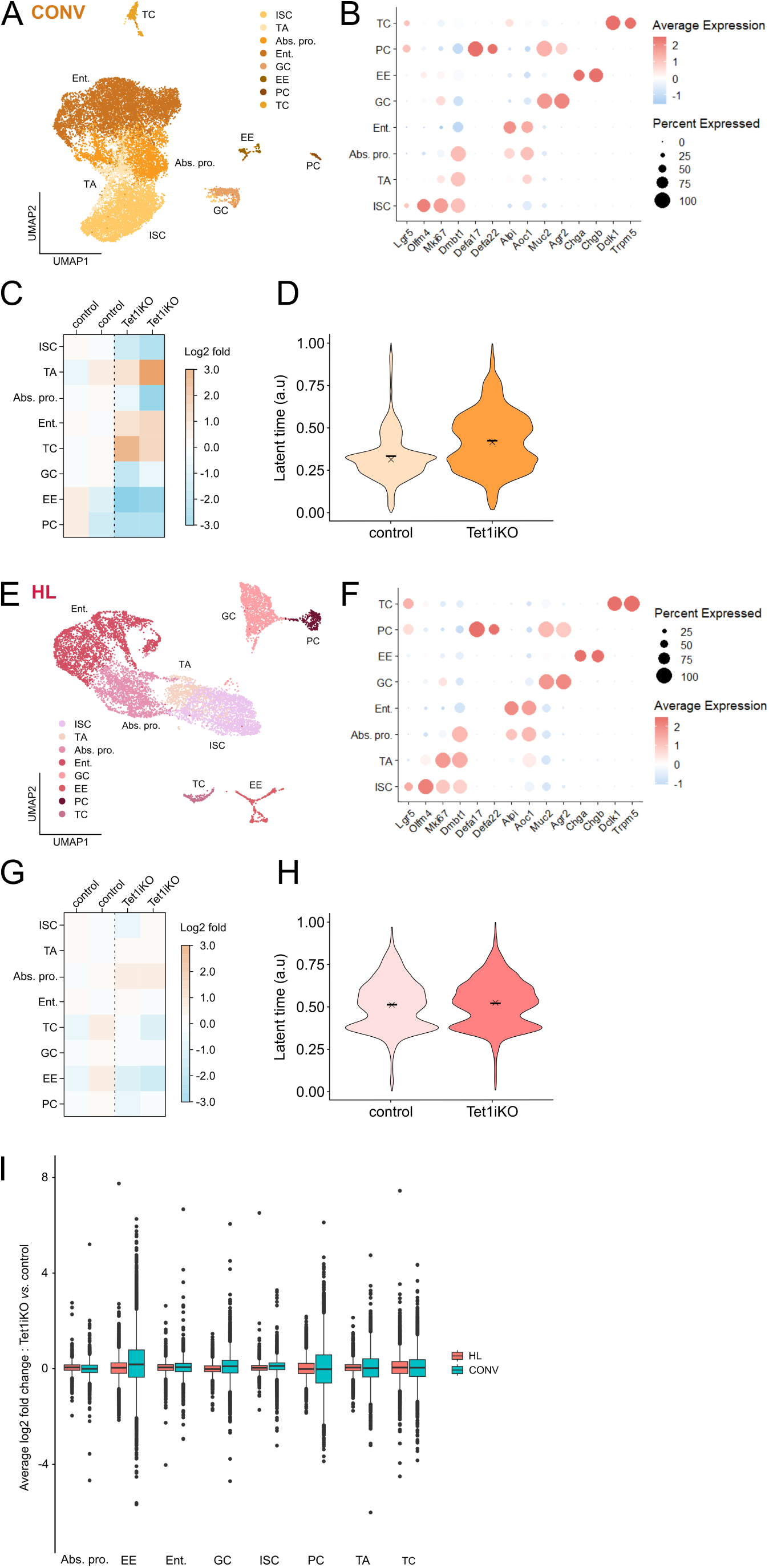
Transcriptomic bias towards absorptive enterocyte and tuft cell differentiation in Tet1iKO intestinal epithelial cells. (A) UMAP of 15,060 single cell transcriptomes from control (n=2) and Tet1iKO (n=2) mice housed in CONV conditions demonstrates expected IEC subpopulations (ISC = Intestinal Stem Cell, TA = Transit-Amplifying cells, Abs. pro. = Absorptive progenitors, Ent. = Enterocytes, GC = Goblet Cell, EE = Enteroendocrine, PC = Paneth Cell, TC = Tuft Cell). (B) IEC marker expression in CONV housing reveals a specific signature for each identified cluster consistent with established IEC subpopulation identity. (C) Log2 fold change (LFC) to analyze cluster enrichment reveals increased proportions of TA, Ent. and TC clusters in Tet1iKO mice from CONV housing relative to controls. All other clusters are proportionally less represented (LFC in cluster enrichment calculated relative to the average control cluster proportion). (D) Latent time values are increased across all IECs from Tet1iKO mice in CONV housing. (E) UMAP and (F) IEC marker expression demonstrates similar clustering results for 10,041 single cell transcriptomes obtained from control (n=2) and Tet1iKO (n=2) mice housed in HL conditions. (G) Relative to CONV housed mice, cluster proportions remain relatively similar between Tet1iKO and control HL mice, with a subtle enrichment in Tet1iKO Abs. pro. (H) Latent time values are also similar between Tet1iKO and control HL mice. (I) LFC between Tet1iKO vs. control samples for each gene in IEC subpopulations reveals a greater magnitude of transcriptomic changes in CONV vs. HL housed samples.

With IEC subpopulations well-defined by scRNA-seq, we next sought to determine cluster proportions between control and Tet1iKO groups for potential lineage allocation bias as observed by histology (Figure S3M and S4M). In CONV housing, we identified a clear positive enrichment for transit-amplifying (TA) progenitor cells and differentiation bias towards enterocyte and tuft cell clusters in Tet1iKO (Figure 2C), consistent with phenotypes identified by histology. Notably, we did not observe the increase in MUC2+ goblet cells identified at the tissue level in single cell transcriptomic data, instead noting decreased goblet cell numbers in Tet1iKO samples (Figure 1B, 2C). This could be due to a higher sensitivity of Tet1iKO goblet cells to lysis during single cell isolation or differences in sampling approaches between histology, which was performed on jejunal sections, and single cell isolation from whole intestine. Interestingly, Tet1iKO mice housed in the CONV facility exhibited decreased ISC numbers and increased TA numbers, leading us to hypothesize that *Tet1*-deficient ISCs might be differentiating faster or prematurely in these conditions (Figure 2C). To test this, we evaluated the overall IEC differentiation rate by implementing latent time analysis. Latent time predicts the relative differentiation stage of individual cells in a data set, based on RNA velocity and splicing ^29^. Tet1iKO IECs from mice housed in the CONV facility exhibited higher latent time values relative to controls, suggesting increased differentiation status across IECs regardless of subpopulation identity (Figure 2D). In HL housing, the same approach was followed. As expected from histological analysis, cluster proportions were much more homogeneous between Tet1iKO and control samples. No notable changes in cell types were observed, except for a subtle enrichment of absorptive progenitors (Abs. pro.) in Tet1iKO (Figure 2G). Latent time values were also unchanged between HL Tet1iKO and controls, suggesting the precocious differentiation phenotype observed in CONV Tet1iKO mice is environmentally dependent (Figure 2H). Our scRNA-seq analyses therefore support observations made by histology and demonstrate that the phenotypic severity of *Tet1* knockout in IECs is dependent on the extrinsic environment.

To further characterize facility-dependent Tet1iKO phenotypes and test the hypothesis that loss of *Tet1* leads to more severe differentiation defects in CONV housing, we conducted differential gene expression analysis. PCA supported a greater overall transcriptomic difference between control vs. Tet1iKO IECs from CONV than control vs. Tet1iKO IECs from HL housed mice (Figure S5A). Additionally, the number of significant differentially expressed genes (DEGs) across all IEC clusters was higher in CONV housing than HL SPF housing (Figure S5B & C). Numbers of DEGs were also higher in each cell type specific cluster in CONV housed Tet1iKO mice (Figure S6, Table S2 & 3). Interestingly, DEGs were greatest among ISC, TA, absorptive progenitors, and enterocyte clusters in both datasets and also most increased in these populations in CONV vs HL mice (Figure S6). To examine gene expression changes in each cell type independent of significance and fold-change cutoffs for DEGs, we plotted the fold change of all genes in Tet1iKO vs. control samples for each IEC cluster. The magnitude of gene expression change in Tet1iKO samples was higher in CONV housing across all clusters (Figure 2I). Together, these analyses support our hypothesis that loss of *Tet1* results in enhanced transcriptomic differences in CONV housing. Importantly, this appears to occur in a largely “lineage agnostic” manner, with all IEC subpopulations exhibiting greater magnitude of gene expression changes in CONV-housed Tet1iKO mice.

### Loss of Tet1 primes intestinal epithelial cells for differentiation at the stem cell level

To further investigate how loss of *Tet1* might result in defective lineage allocation, we sub-clustered ISCs from scRNA-seq data. In both data sets, sample and group distribution were rather homogeneous except for one Tet1iKO and one control sample from CONV housing exhibiting non-overlapping cells by UMAP (Figure S7A & B and F & G). We selected 41 PCs and a resolution of 0.3, leading to the identification of 4 ISC clusters in CONV housing (Figure S7 C-E and Figure 3A). 42 PCs and a resolution of 0.3 were selected in HL housing, also leading to the identification of 4 ISC clusters (Figure S7H-J and Figure 3J).

**Figure 3.**
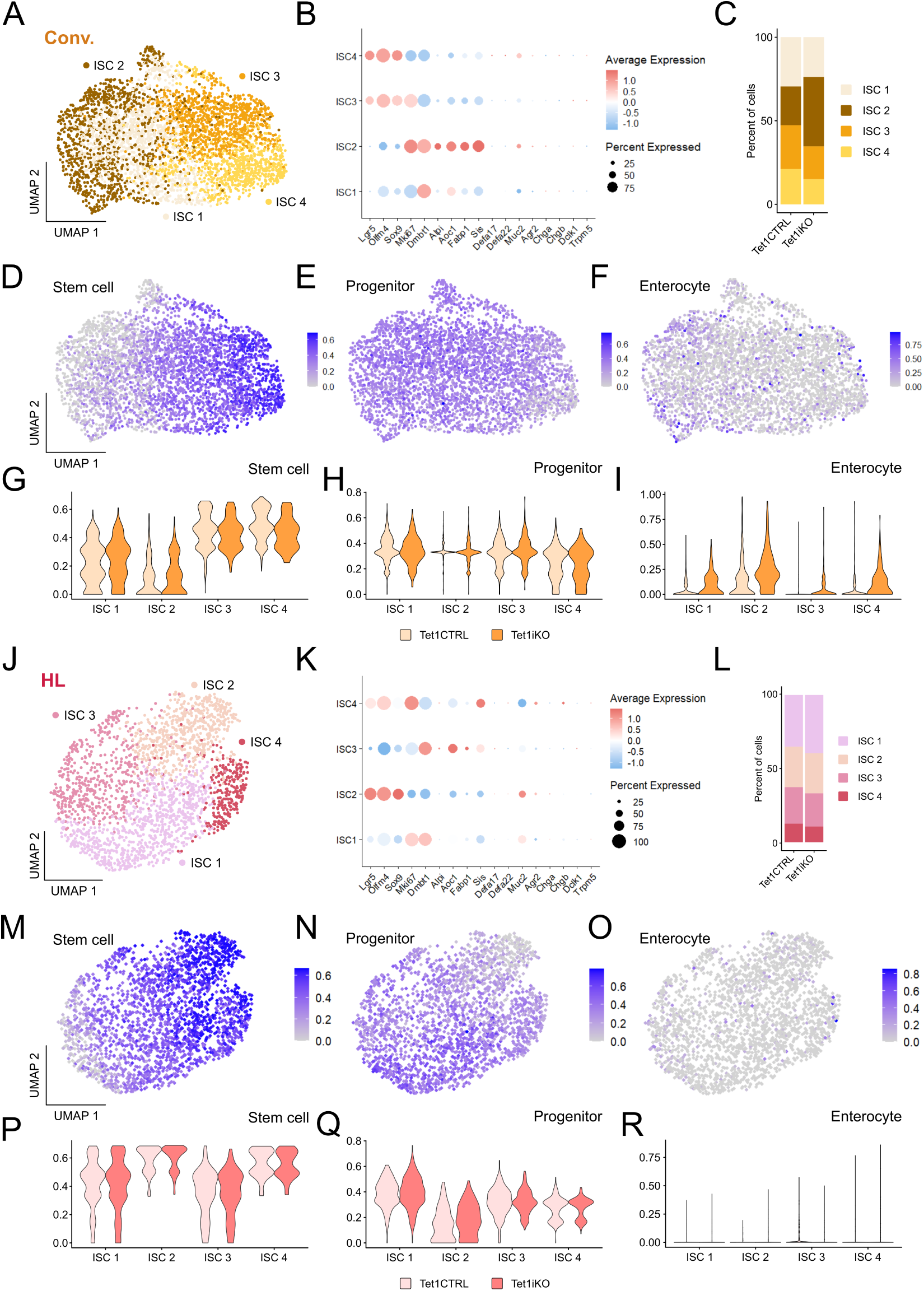
Tet1iKO ISCs upregulate absorptive enterocyte genes in conventional housing. (A) scRNA-seq from CONV housed mice identifies 4 ISC subclusters. (B) IEC marker enrichment reveals higher expression of ISC markers *Lgr5, Olmf4* and *Sox9* in ISC3 and ISC4. ISC2 is enriched for *Mki67, Dmbt1, Alpi, Aoc1, Fabp1*, *Sis* and *Muc2*. ISC1 expresses TA-associated *Dmbt1* over other markers. (C) Cluster proportion analysis reveals that a larger number of ISCs in Tet1iKO mice belong to ISC2. In CONV housing, UMAP for UCell gene signatures shows (D) enrichment for stem cell genes in ISC3 and ISC4, (E) homogenous expression of progenitor genes across ISC subclusters, and (F) a subtle enrichment of enterocyte genes in ISC2. (G) UCell gene signature enrichment is unchanged between Tet1iKO and control across all ISC subclusters for (G) stem cell genes and (H) progenitor genes, but (I) the enterocyte signature is enriched in Tet1iKO compared to control mice across all ISC clusters. (J) 4 ISC subclusters are also identified in HL housed mice. (K) ISC markers *Lgr5, Olmf4, Sox9* and *Muc2* are most enriched in ISC2, while ISC1 is enriched for *Mki67, Dmbt1* and *Muc2* expression. ISC3 is enriched for *Dmbt1, Aoc1, Fabp1* and *Sis*. ISC4 expresses *Lgr5, Sox9* and is enriched for *Mki67* and *Sis* expression. (L) HL ISC subpopulation proportions are highly similar between Tet1iKO and control. (M) UMAP for UCell gene signatures shows enrichment for the stem cell signature over all 4 HL ISC subclusters, but predominantly in ISC2 and ISC4. (N) The progenitor signature is also detected broadly across all subclusters and enriched in ISC1, while (O) the enterocyte signature is not reliably detected in any of the HL ISC subclusters. Comparing differences in UCell gene signature scores shows no change between HL Tet1iKO and control ISCs for (P) stem cell, (Q) progenitor, or (R) enterocyte gene signatures.

We first applied a defined IEC gene set (Table S1) to deduce the relative identity of each ISC cluster by assessing the expression of a restricted list of canonical stem, progenitor, and mature IEC markers (Figure 3 B & K). In CONV housing, ISC clusters 3 and 4 are most enriched for ISC genes. Cluster 2 has the highest enrichment for TA and differentiated genes. Cluster 1 exhibits an intermediate profile oriented towards the expression of progenitor genes, but not yet expressing differentiated transcripts (Figure 3B). Interestingly, quantification of cluster proportions revealed an enrichment for ISC cluster 2 in Tet1iKO at the expense of ISC clusters 1, 3 and 4, suggesting a CONV Tet1iKO ISC pool “skewed” toward differentiation (Figure 3C). To more accurately describe ISC cluster identity and enrichment between groups, we conducted gene set enrichment analysis using UCell signature estimation for stem cell, progenitor and absorptive enterocyte signatures. Specific gene sets were derived from previously published intestinal scRNA-seq (Table S1) ^28^. The stem cell signature was most enriched in clusters 3 and 4 and relatively unchanged between Tet1iKO and control samples (Figure 3D & G). Similarly, the TA progenitor gene set was homogenously represented across ISC clusters and evenly enriched between Tet1iKO and control groups (Figure 3E & H). Interestingly, the absorptive enterocyte gene set was most enriched in ISC cluster 2 among control subpopulations, but universally upregulated in all Tet1iKO subpopulations vs. all control subpopulations (Figure 3F & I). Together, ISC subpopulation analysis of CONV housed mice shows that Tet1iKO ISCs are skewed toward an early differentiation phenotype, enriched for absorptive enterocyte gene signatures.

In HL housing conditions, ISC cluster 2 exhibited the highest enrichment for stem cell genes (Figure 3K). The other 3 clusters had intermediate expression profiles between stem cells, progenitor and enterocyte-associated genes (Figure 3K). Unlike in CONV ISC subpopulations, we did not observe differences in cluster proportions between Tet1iKO and control ISCs from the HL facility (Figure 3L). Similarly, gene set enrichment analysis did not reveal major differences between ISC subpopulations or Tet1iKO and control groups (Figure 3 M-R). Stem cell and TA progenitor signatures were both homogenously expressed across ISC subpopulations (Figure 3M & P and 3N & Q). Unlike in CONV ISC subpopulations, the absorptive enterocyte signature was almost non-existent across all HL ISC clusters, regardless of control or Tet1iKO genotype (Figure 3 O & R). Overall, our ISC-specific single cell transcriptomic analysis reinforces observations in full IEC scRNA-seq datasets that absorptive enterocyte differentiation is enhanced in CONV-housed mice. Compellingly, this phenotype is apparent when comparing CONV vs. HL control ISCs, and is more pronounced in Tet1iKO. Further, enhanced absorptive enterocyte specification following loss of *Tet1* appears to occur at the ISC level. Importantly, the lack of premature absorptive specification in HL Tet1iKO ISCs suggests that *Tet1*-associated phenotypes only arise in response to an external stimulus.

### Loss of Tet1 sensitizes intestinal epithelial cells to extrinsic signaling

Our *in vivo* data suggest that *Tet1* regulates ISC responsiveness to changes in extrinsic signaling, such as those associated with different animal housing conditions and *Tmu* colonization status. Intriguingly, HL housing appears to “silence” the Tet1iKO phenotypes apparent in CONV housing. Because *in* vivo housing conditions consist of many confounding and difficult to define variables, we reasoned that we could leverage the lack of HL phenotype to test the responsiveness of Tet1iKO IECs to a range of controlled stimuli. First, we generated organoids from Tet1iKO and control intestinal crypts and stimulated them with the Notch inhibitor DAPT or IL-4 to induce broad secretory differentiation or tuft cell hyperplasia, respectively ^30,31^. Numbers of DCLK1+ tuft cells were significantly increased in Tet1iKO organoids in both DAPT and IL-4 media conditions, consistent with reports that both Notch inhibition and type II cytokines can stimulate tuft cell hyperplasia (Figure 4A & B) ^32^. As expected, IL-4 treatment had no effect on CHGA+ enteroendocrine numbers, but DAPT-treated organoids exhibited increased enteroendocrine cells that were significantly higher in Tet1iKO samples (Figure 4C & D).

**Figure 4.**
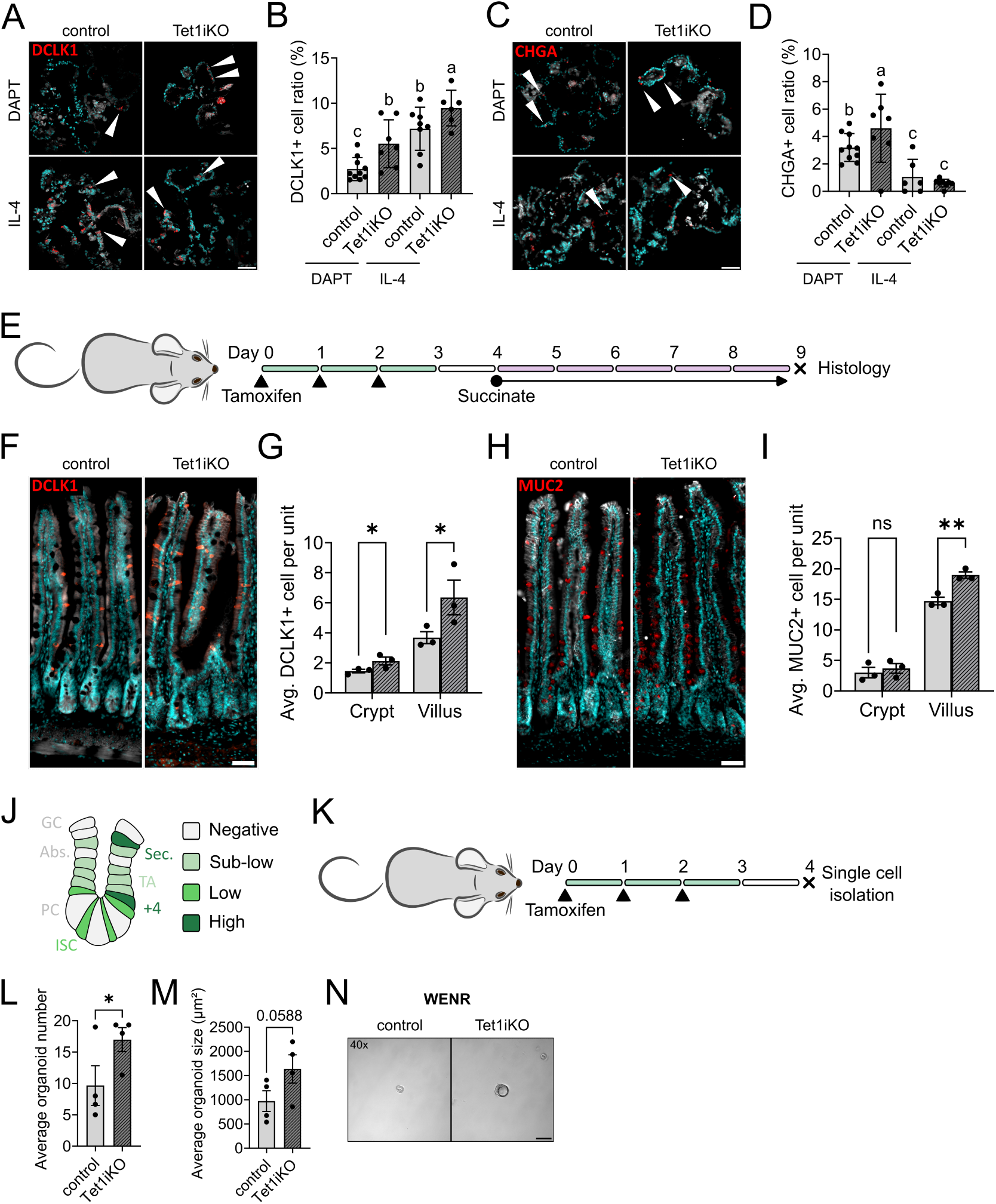
Tet1iKO enhances intestinal epithelial response to extrinsic signaling in vitro and in vivo. (A, B) DCLK1+ tuft cells are significantly increased in Tet1iKO organoids subjected to 48h in differentiation media containing either DAPT or IL-4 following 4 days of establishment and expansion (white arrow heads indicate DCLK1+ cells). (C, D) CHGA+ enteroendocrine cells are increased in Tet1iKO organoids subjected to DAPT, but not IL-4 (white arrow heads indicate CHGA+ cells) (scale bars represent 50 µm) (each group represents n = 6 to 10 individual organoids, different letters indicate statistically significant differences between groups, p < 0.05 one-way ANOVA). (E) To induce tuft and goblet cell hyperplasia *in vivo*, control and Tet1iKO mice were subjected to tamoxifen recombination, then administered succinate *ad libitum* in drinking water for 5 days prior to tissue collection. (F, G) Succinate treated Tet1iKO mice exhibited a significantly greater increase in tuft cell number relative to controls in both crypts and villi. (H, I) Goblet cell hyperplasia was also more pronounced in Tet1iKO mice, but restricted to villi (scale bars represent 100 µm) (control n = 3, Tet1iKO n = 3,* for p < 0.05 and ** for p < 0.01). (J) Schematic representation of *Sox9^EGFP^* expression in intestinal crypts, highlighting differential EGFP expression levels associated with distinct IEC populations. (K) For single cell organoid-forming assays, HL housed Tet1iKO and control mice carrying a *Sox9^EGFP^* reporter transgene were subjected to tamoxifen recombination and *Sox9^low^* ISCs were isolated by FACS (L) Tet1iKO ISCs form significantly greater numbers of organoids after 5 days of culture in WENR media. (M) Tet1iKO organoid size is not significantly larger than controls, but trends toward significance (control n = 4, Tet1iKO n = 4 independent biological replicates; * indicates p < 0.05). (N) Representative brightfield images of organoids after 5 days of culture, demonstrating increased size of Tet1iKO organoids (scale bar represents 50 µm).

Next, we wanted to determine if Tet1iKO IECs would exhibit enhanced differentiation potential *in vivo*, again focusing on secretory lineages due to their ease of quantification by histology and involvement in our initial Tet1iKO phenotype. Protists produce succinate, which is detected by tuft cells and subsequently activates a type II immune response, resulting in tuft and goblet cell hyperplasia ^23^. Administration of succinate in drinking water reproduces this effect ^23,31^. We induced *Tet1* recombination as previously described, then administered succinate *ad libitum* for 5 days before collecting tissue for histology (Figure 4E). Compellingly, Tet1iKO intestines exhibited significantly higher numbers of villus-localized tuft and goblet cells (Figure 4F-I). Together with *in vitro* organoid experiments, these data demonstrate that Tet1iKO IECs are more sensitive to differentiation, even in the absence of a baseline phenotype.

Our *in vivo* and *in vitro* data demonstrate increased absorptive and secretory differentiation sensitivity in Tet1iKO intestines, respectively. Elevated OLFM4+ IEC numbers in our initial Tet1iKO phenotype prompted us to ask if loss of *Tet1* also increases sensitivity to pro-ISC signaling. To isolate ISCs, we generated Tet1iKO and controls carrying a Sox9^EGFP^ BAC transgenic reporter allele. Sox9^EGFP^ is expressed at distinct levels and has been used by our lab and others to isolate differentiated IECs (*Sox9^neg^*), TAs (*Sox9^sublow^*), ISCs (*Sox9^low^*), and enteroendocrine cells/tuft cells/secretory progenitors (*Sox9^high^*) (Figure 4J) ^8,33–35^. *Tet1* recombination was induced as previously described, and single *Sox9^low^* ISCs isolated by FACS 2 days after the last dose of tamoxifen (Figure 4K). Tet1iKO and control ISCs were cultured in high-WNT3A containing culture conditions previously reported to promote organoid formation ^36^. Strikingly, Tet1iKO ISCs formed significantly higher numbers of organoids, which trended toward being larger in size (Figure 4L-N). These data demonstrate that Tet1iKO ISCs are more functionally sensitive to WNT ligands than controls, exhibiting increased stemness through enhanced organoid formation. Together with the observed response to pro-differentiation signaling, these experiments suggest that *Tet1* serves as a general “buffer” that limits the magnitude of ISC response to extrinsic stimuli.

### Tet1 modulates transcription factor binding without impacting ISC chromatin accessibility

*Tet1* exerts broad regulatory influence on the genome, functioning via catalytically dependent and independent mechanisms that involve DNA methylation and co-recruitment of other chromatin regulatory complexes ^37^. We reasoned that assaying chromatin accessibility in HL housed mice would provide an overall snapshot of how *Tet1* impacts the chromatin landscape in the absence of specific, phenotype-inducing signaling. Tet1iKO and control ISCs were isolated from Sox9^EGFP^ mice and subjected to ATAC-seq ^38^. PCA of aligned reads did not reveal distinct clustering of Tet1iKO and control ISCs, demonstrating no significant variance by genotype (Figure 5A). DiffBind analysis to identify differentially accessible chromatin supported PCA, detecting only 2 and 3 peaks that were more and less accessible in Tet1iKO ISCs, respectively (Figure 5B, Table S4). Visualizing ATAC-seq data in browser tracks revealed reproducible peaks with strong signal:noise ratio that remained unchanged, even surrounding *Lars2* and *Dnajb1*, two genes that were differentially expressed by scRNA-seq in ISCs from both CONV and HF housed Tet1iKO mice (Figure S5B & C and 5C).

**Figure 5.**
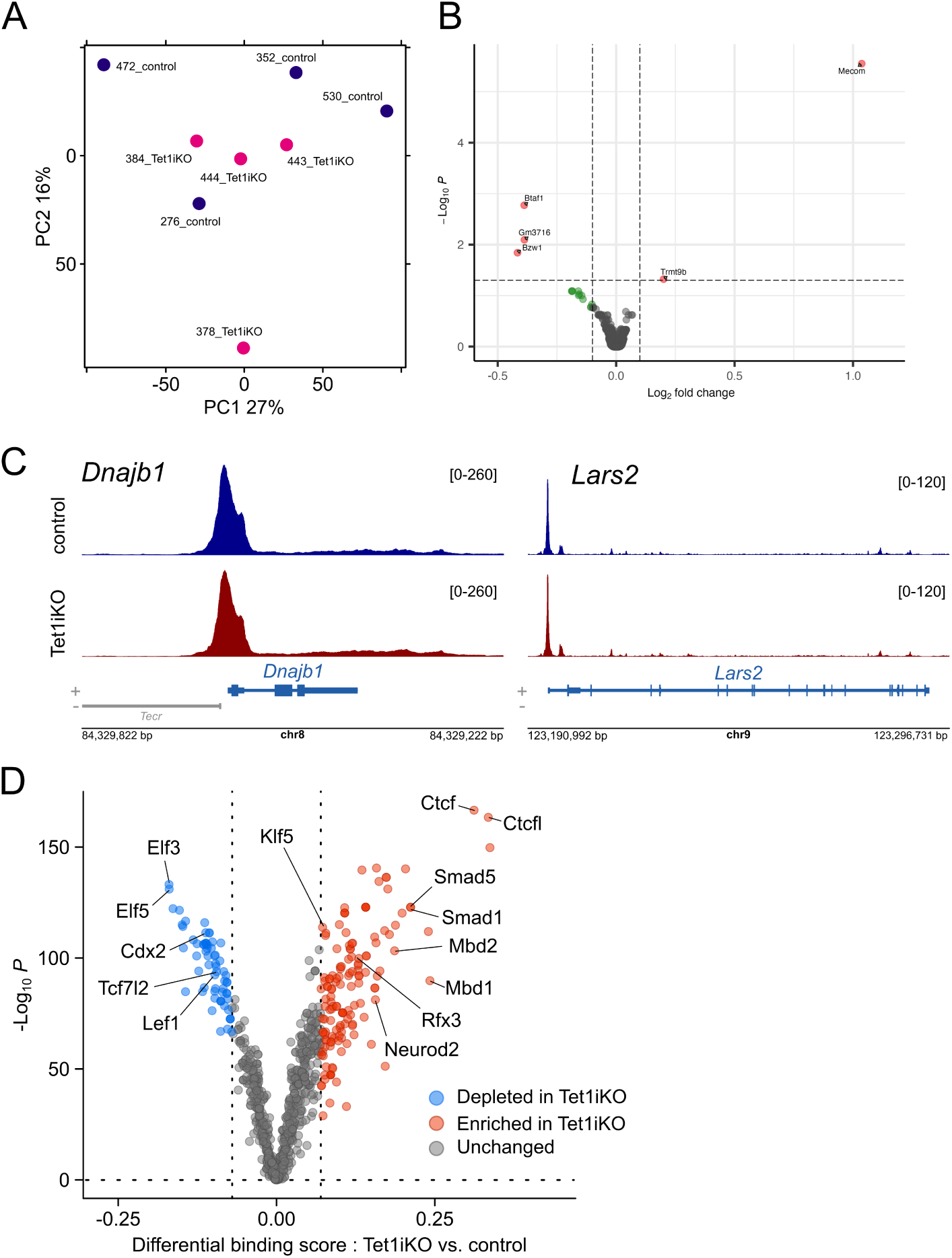
Loss of Tet1 has minimal impacts on ISC chromatin accessibility but affects TF binding. (A) PCA of ATAC-seq from Sox9^low^ ISCs demonstrates a lack of global differences in chromatin accessibility between Tet1iKO and control samples in HL housing. (B) DiffBind analysis confirms a minimal change in chromatin accessibility following loss of *Tet1* (significant peaks labeled by nearest gene). (C) ATAC-seq signal surrounding *Dnajb1* and *Lars2*, which are significantly upregulated in Tet1iKO mice regardless of housing condition, identifies reproducible peaks but no differences between control and Tet1iKO mice (browser tracks represent merged ATAC-seq signal from n = 3 control and n = 3 Tet1iKO). (D) TOBIAS footprinting analysis predicts significant changes in TF binding in Tet1iKO samples, including increased binding of pro-differentiation TFs and decreased binding of ISC-associated TFs.

Some degree of IEC plasticity and lineage allocation is known to be driven independent of changes in chromatin accessibility, consisting instead of TFs acting on a “broadly permissive” chromatin landscape ^4,5^. To this end, we analyzed potential TF binding using TOBIAS, a computational footprinting pipeline for ATAC-seq data ^39^. Despite the lack of changes in chromatin accessibility, TOBIAS predicted increased binding of 54 TFs and decreased binding of 44 TFs in Tet1iKO ISCs (Figure 5D). Methyl-binding domain TFs, MBD1 and MBD2, were predicted to have increased binding following loss of *Tet1*, consistent with increased DNA methylation which would be expected with reduced TET1 conversion of 5mC to 5hmC ^12^. Differentiation related TFs predicted to have increased binding in Tet1iKO ISCs included SMAD1 and SMAD5, which respond to BMP signaling and are broadly implicated in ISC differentiation, as well as NEUROD2 and RFX3, which are involved in enteroendocrine specification ^40–42^. Interestingly, core ISC regulatory TFs LEF1 and TCF7L2 (also known as TCF4) were predicted to have decreased binding in Tet1iKO ISCs (Figure 5D). Several other TFs broadly implicated in IEC development and ISC differentiation were also predicted to have dysregulated binding in Tet1iKO ISCs, including multiple CDX, KLF, and ELF family TFs (Figure 5D and Table S5). Finally, CTCF and CTCFL, which are involved in higher order chromatin organization and long-range enhancer-promoter interaction, were predicted to exhibit the largest increase in binding in Tet1iKO ISCs (Figure 5D)^43^. Taken together, our ATAC-seq analyses suggest that *Tet1* does not directly impact chromatin accessibility but may modulate the ability of TFs to bind target loci in ISCs. Compellingly, the panel of TFs predicted to have increased and decreased binding following loss of *Tet1* fit a profile of increased differentiation potential with attenuated stemness.

## Discussion

The relevance of the *cis*-regulatory chromatin landscape in ISC biology has remained somewhat controversial, with some studies suggesting that TF regulation is ultimately more important for driving cell fate decisions ^4,5^. In this study, we show that *Tet1* acts broadly to “fine tune” how IECs respond to extrinsic signaling. We identify several phenotypes *in vivo* that are dependent on the housing conditions of Tet1iKO and control mice and do not follow easily explained patterns typically associated with TF regulators of cell fate. Specifically, defects observed in Tet1iKO mice are not restricted to one or even several closely related IEC subpopulations, but rather involve both absorptive and secretory lineages. These data highlight the intrinsic challenges in studying *Tet1* in ISCs and may be relevant to other chromatin modifying enzymes with similarly broad regulatory impacts. The ability to identify housing conditions that “silenced” phenotypic differences in Tet1iKO mice was ultimately critical in providing a baseline to challenge the relative responsiveness of Tet1iKO ISCs to different signaling environments.

Our results differ significantly from previous studies using *Tet1*-null mice. These described a significant decrease in ISC numbers as well as reduced WNT signaling, which is required to maintain the *Lgr5*+ ISC pool ^10^. While our data support precocious differentiation of Tet1iKO ISCs at the transcriptomic level *in vivo*, we also show that isolated Tet1iKO ISCs are more functionally responsive to high WNT signaling *in vitro*. In this regard, our single cell organoid forming experiments suggest that loss of *Tet1* in adult ISCs does not simply promote differentiation, but rather increases sensitivity to pro-differentiation signals. As ISCs are relatively rare and constantly competing for access to niche signals, it is likely that the increased tendency toward differentiation of Tet1iKO ISCs *in vivo* is simply a response to the dominant signaling environment ^44,45^. One compelling explanation for why our results may differ from those in *Tet1*-null intestines may be related to the role of TET1 in DNA methylation. MBD pulldown experiments, which allow for mapping of 5mC via affinity purification, demonstrated that most adult ISC-related genes are methylated in embryonic IECs ^9^. This suggests that a wave of demethylation is associated with intestinal maturation. It is possible that this process is *Tet1*-dependent, especially given the critical role of TET1 in similar broad demethylation essential for germline reprogramming and induced pluripotency ^46,47^. This could result in a developmental-specific role for *Tet1* in establishing the *Lgr5* ISC pool independent of its role in adult ISC differentiation, which involves fewer changes in DNA methylation ^8,9^.

Taken together, our data support a model where *Tet1* “buffers” the responsiveness of ISCs to the extrinsic environment. We propose a model in which TET1 is a modulator – not a driver – of ISC fate, acting to prevent premature binding of TFs induced by upstream signals, but not independently sufficient to drive or prevent differentiation. Our findings are somewhat similar to recent reports that H3K36me3 also has subtle but significant impacts on IEC fate ^34^. In these studies, a dominant-negative H3K36M mutant allele was used to block methylation at H3K36. This resulted in a subtle dysregulation of gene expression in mature IEC lineages, associated with ultrastructural changes in some secretory cell types ^48^. Unlike our results, H3K36M mice also activated damage-associated facultative stem cell transcriptomic programs in the absence of injury ^48^. However, both examples suggest that chromatin in the intestine acts to reinforce and preserve cell identities, raising or lowering barriers to changes in cell fate.

Results from TF footprinting analysis in ISCs of HL housed Tet1iKO mice are consistent with increased differentiation and decreased stemness, and further support a general model where *Tet1* inhibits TF binding. Interestingly, our scRNA-seq results from the same facility do not demonstrate any detectable transcriptomic changes consistent with decreased ISC numbers or premature differentiation. One limitation of our study is that we were unable to identify a specific genetic regulatory mechanism explaining how the loss of *Tet1* increases ISC sensitivity to extrinsic signals. Because TET1 interacts with a range of chromatin regulatory complexes, including HDACs and PRC2, and participates in DNA demethylation, it is likely that several different mechanisms may contribute to this phenotype. Future studies to profile the chromatin landscape of Tet1iKO ISCs will be needed to dissect these mechanisms.

## Material & methods

### Mice models & housing conditions

All experiments were carried out using mice between 8 and 24 weeks of age, maintained on C57Bl/6 background. Previously described *Sox9^EGFP^* and *Vil-CreER* mice have been obtained from Magness lab (UNC, Chapel Hill, NC, USA) ^16,49^. Previously described *Tet1^fl/fl^* mice were obtained from Rao lab (LJI, San Diego, CA, USA) ^17^. All transgenic/mutant alleles were maintained at heterozygosity, except *Tet1^fl/fl^* maintained at homozygosity. Knockout efficiency was confirmed by RT-qPCR for *Tet1* expression relative to control animals from isolated ISCs (Figure S1).

Animals received PicoLab Rodent Diet 20 (LabDiet, 5053) and water *ad libitum*. Housing has been maintained under specific pathogen free (SPF) status over three different facilities throughout the study, although presenting different biological surveillance levels. Conventional (CONV) SPF housing was maintained at the University of North Carolina (Chapel Hill, NC, USA) and at Whitehead Biomedical Research Building (WBRB) Emory University (Atlanta, GA, USA). High-level (HL) SPF housing was maintained at the Health Sciences Research Building (HSRB) at Emory University (Atlanta, GA, USA). HL SPF includes additional surveillance to CONV SPF housing for organisms relevant to our study such as *Tritrichomonas muris*. The University of North Carolina and Emory University Institutional Animal Care and Use Committee reviewed and approved all animal protocols.

### Tamoxifen induced Tet1 knockout

*Tet1^fl/fl^:VilCreER* (Tet1iKO) animals received daily intraperitoneal injection of 100 µl of 10 mg/ml tamoxifen (Sigma Aldrich T5648) for 72h to ensure maximum penetrance of Cre-mediated recombination. Tamoxifen was freshly prepared at the beginning of treatment protocol. After resuspending tamoxifen in 10% Ethanol:90% sunflower oil, suspension was sonicated 3 times for 30 seconds with 1 minute intervals on ice. Tissue was harvested 96h hours after the first injection allowing a 48h tamoxifen clearance and penetrance of the deletion through tissue renewal. To anticipate for Cre recombinase off-target and tamoxifen side effects, *VilCreER* animals were employed as control.

### Succinate water treatment

Succinate water was prepared at 100 mM of succinic acid in 500 ml water bottle. pH was adjusted to 5.5 with NaOH and water sterilized by vacuum filtration through a 0.2 µm PES filter. Water was provided *ad libitum* after tamoxifen protocol.

### T muris screening

Cecal tissue was harvested, opened longitudinally and dropped in 10 ml PBS in a 50 ml conical tube. After content homogenization by hand shaking, volume was brought up to 50 ml with PBS. 10 µl of suspension were mounted on microscopic slide and coverslip then observed under microscope in brightfield for *T. muris* colonization. Representative pictures were acquired on Olympus IX-83 inverted epifluorescent microscope and CellSens Imaging software at 20x magnification.

### Intestinal crypt isolation and cell dissociation

Crypts were isolated for organoid culture as previously described by Zwarycz et al.^50^ Briefly, intestines were dissected out, the first 6 cm discarded, and the remaining proximal half designated as jejunum and taken for organoid isolation. Jejunal segments were opened longitudinally and rinsed briefly in a 50 ml conical containing 10 ml Dulbecco’s Phosphate-Buffered Saline (DPBS, Gibco 14190250). Tissue was transferred to a new 50 ml conical containing 3 mM EDTA (Corning 46-03-Cl) in 10mL sterile DPBS and incubated at 4°C on a rocking platform set to 25 RPM for 15 minutes. Tissue was retrieved and villi removed by gentle “brushing” with a P200 pipette tip on a glass plate. Intestinal tissue was then rinsed briefly in a petri dish containing sterile DPBS and cut into pieces ∼0.5 cm in length before being transferred to a new 50 ml conical containing 3 mM EDTA in 10 ml sterile DPBS and incubated at 4°C on a rocking platform for 35 minutes. Jejunal pieces were transferred to a 50 ml conical containing 10 ml sterile DPBS and shaken for 3 to 4 minutes to release crypts, confirming expected crypt density and morphology by light microscopy at 1 minute intervals to determine when to end the dissociation protocol. 10 ml sterile DPBS was added to isolated crypts, which were then filtered through a 70 µm cell strainer (Falcon 352350), pelleted at 600g for 5 minutes at 4°C, and resuspended in 200-500 µl Advanced DMEM/F12 (Gibco 12634010).

For single-cell dissociation, released intestinal crypts were not filtered but centrifuged at 600g for 5 minutes at 4°C. After discarding supernatant, pellet was resuspended in 20 ml HBSS with 0.3 U/ml Dispase (Corning 354235) and 0.2 mg/ml DNase (Sigma Aldrich), and incubated at 37°C for 12 to 16 minutes with gentle shaking and observation by light microscopy every 2 minutes until approximately 80% of the suspension consisted of single cells. Reaction was quenched with 30 ml HBSS and suspension filtered with a 40 µm cell strainer (Falcon 352340). Filtered suspension was centrifuged at 600g for 5 minutes at 4°C and supernatant discarded. Pellet was resuspended in 10 ml ice cold DPBS and suspension centrifuged at 600 g for 5 minutes at 4°C. After discarding supernatant, pellet was resuspended in 1 ml sorting media : Advanced DMEM/F12 (Gibco 12634028), 1X N2 (Gibco 17502048), 1X B27 w/o vitamin A (Gibco 12587010), 1X HEPES (Gibco 15630080), 1X Penicillin/Streptomycin (Sigma-Aldrich P4333-100ML), 1X Glutamax (Gibco 35050061). Suspension was further processed for FACS enrichment.

### FACS

Dissociated single cells in sorting media were stained for surface markers 1:100 CD44-BV421 (Biolegend 103039), 1:100 CD326-APC/Cy7 (Biolegend 118218), 1:100 CD31-APC (Biolegend 102510) and 1:100 CD45-APC (Biolegend 103112) for 45 minutes on ice. Stained cells were rinsed once with Advanced DMEM/F12, centrifuged at 600g for 5 minutes at 4°C and resuspended in sorting media. 5 µl of 7-AAD (Biolegend 420404) and 5 µl of Annexin V-APC (Biolegend 640941) were added 15 minutes before sorting. Cells were sorted on Sony SH800 FACS instrument with a 100 µm disposable nozzle chip. For single-cell RNA sequencing gating was carried out as described in Fig. S8. Debris were excluded based on size via bivariate plot of FSC-A against BSC-A. Doublets/multimers were excluded using bivariate plot of FSC-H against FSC-A and BSC-H against BSC-A. Dead cells and non-epithelial cells were excluded based on CD45 negative/CD31 negative/7AAD negative/Annexin V negative. CD326 and CD44 stainings provided indication for CD326 positive epithelial cells enrichment and CD44 positive cryptic cell ratio versus CD44 negative non-cryptic cells. For *Sox9^low^* intestinal stem cell enrichment, additional gating for Sox9^EGFP^ populations was carried out as described previously and shown in Figure S9 ^8^.

### Organoid culture

Crypt density per 10 µl media was examined qualitatively by light microscopy and crypt density estimated, with the goal of determining volume required to achieve between 50-100 crypts per 10 µl Matrigel in culture. Crypts were resuspended in 75% phenol red-free, growth factor-reduced Matrigel (Corning 356231) and plated as 10 ml droplets in 96 well plates or 40 ml droplets in 48 well plates. Matrigel was allowed to polymerize at RT for 1 min, then at 37°C for 20 minutes. Media was overlaid at 100 µl per well in 96 well plate and 200 µl per well in 48 well plates. Basal ENR media was composed of [Advanced DMEM/F12 (Gibco 12634028), 1X N2 (Gibco 17502048), 1X B27 w/o vitamin A (Gibco 12587010), 1X HEPES (Gibco 15630080), 1X Penicillin/Streptomycin (Sigma-Aldrich P4333-100ML), 1X Glutamax (Gibco 35050061), 10% RSPO1-CM (made using RSPO1 transfected HEK293T cells following manufacturer protocol: Sigma-Aldrich SCC111), 50 ng/ml recombinant murine EGF (Fisher Scientific PMG8041), and 100 ng/ml recombinant human Noggin (Preprotech 120-10C-20UG)]. 500 µg/ml Primocin (Invivogen ant-pm-1) and 10 mM Y27632 (Selleck Chemicals S1049) were added to overlay media for the first 48h after plating crypts and then excluded from all other media changes. For promotion of stemness, ENR media was supplemented with 40% WNT3a-CM (WENR) (made using Wnt3a transfected L-M(TK-) cells following manufacturer protocol : ATCC CRL-2647). For secretory differentiation purposed, ENR media was supplemented with 25 µM Notch inhibitor DAPT (SelleckChem S2634). For tuft cell differentiation induction, ENR media was supplemented with type 2 cytokine IL-4 100 ng/ml (Biolegend 574304).

### Cytohistology

After tissue harvest, small intestines were flushed with DPBS, fixed overnight at 4°C with 4% paraformaldehyde (PFA, Thermo Fisher 41678-5000) in DPBS then transferred to 30% sucrose (Fisher bioreagents BP220-10) in distilled water for 24h at 4°C. Small intestines were then opened longitudinally and embedded in Tissue-Plus OCT (Fisher Scientific 23-730-571) following the swiss-roll method. Embedded tissues were frozen on dry ice before storage at -70°C.

Organoid fixation and embedding were performed following as previously described ^51^. Procedure required pre-coated pipet tips and conical tubes with 1% BSA (Sigma-Aldrich A9647) in DPBS to limit loss of material. Intestinal organoids were retrieved with Cell Recovery Solution (Corning 354253) in a 1.7 ml conical tube and incubated at 4°C for 45 minutes. After Matrigel dissociation by gentle pipetting, organoids were pelleted at 100 g for 1 minute. Supernatant was gently discarded and organoids fixed with 4% PFA in DPBS for 20 minutes on ice. Organoids were pelleted at 100 g for 1 minute and supernatant discarded. Pellet was gently washed with DPBS and centrifuged at 100 g for 1 minute. Supernatant was discarded and organoids resuspended with 2% Methylene Blue (Cayman chemical company 26095) in DPBS for 20 minutes at room temperature (RT). Organoids were pelleted at 100 g for 1 minute and supernatant discarded. Pellet was gently washed in DPBS followed by centrifugation at 100 g for 1 minute and supernatant discarded. This washing step was repeated 3 times until supernatant was clear. Organoids were then resuspended in 30% sucrose in distilled water overnight at 4°C. After centrifugation at 100 g for 1 minute, supernatant was discarded. Organoids were embedded in OCT and frozen on dry ice before storage at -70°C.

Cytohistology blocks were sectioned on a cryostat (Leica CM1860) at 10 µm intervals. Sections were retrieved on charged microscopy slides (MED-Vet international 9308W) and allowed to dry for 30 minutes to 1h at RT. Slides were stored at -70°C.

### Immunofluorescence

Sections were washed 3 times with Phosphate-Buffered Saline (PBS, Corning 46-013-CM). Antigen retrieval was performed if recommended by supplier. Slides were immerged in Reveal Decloaker RTU solution (Biocare Medical RV1000MMRTU), brought to 114-121°C at high pressure for 1 minute in pressure cooker and brought back to RT. Slides were then permeabilized for 20 minutes with 0.3% Triton X-100 (Sigma T8787-250ML) in PBS at RT and blocked in 5% normal donkey serum (NDS, Jackson Immuno 017-000-121) for 45 minutes at RT. Primary antibodies were diluted in PBS and incubated at 4°C overnight. Secondary antibodies were diluted in PBS and incubated for 45 minutes at RT. Nuclear counterstain was carried out using DAPI (Millipore Sigma; D9542) diluted 1:2000 in PBS and incubated 10 minutes at RT. Primary antibodies were used at the following concentrations for immunostaining: 1:200 anti-CD326 (EPCAM, BioLegend 118201), 1:100 anti-DCAMKL1 (DCLK1, Abcam ab192980), 1:250 anti-CHGA (Immunostar 20085), 1/25 anti-FABP1 (Invitrogen PA5-47159), 1/100 anti-LYSOZYME (LYZ, Diagnostic Biosystems RP028), 1:250 anti-MUCIN2 (MUC2, Santa Cruz sc-15334), 1:200 anti-OLFM4 (Cell Signaling 39141S). Secondary antibodies used for immunostaining: 1:500 donkey anti-rabbit Alexa Fluor 555 (ThermoFisher A31572), 1:500 donkey anti-rat Alexa Fluor 488 (ThermoFisher A21208). Slides were mounted with homemade Mowiol mounting media and cured overnight in the dark at RT before storage at 4°C.

### Imaging and quantification

Representative images were acquired on an inverted Olympus FV1000 confocal microscope system and Olympus Fluoview v4.2 software in the Emory Integrated Cellular Imaging Core. Images for quantification were taken using an Olympus IX-83 inverted epifluorescent microscope and cellSens Imaging software. For tissue sections, 50 crypts and/or villi were acquired per biological replicate at 20x magnification. For organoids, up to 10 fields were acquired per experimental group at 40x magnification.

Quantification of positive cells on tissue sections was performed during acquisition by eye or post-acquisition by hand with ImageJ Fiji software. Quantification of nuclei on organoid sections was automated with a macro made with ImageJ Fiji software. Macro pipeline includes DAPI channel extraction, automatic contrast adjustment, Gaussian blurring and automatic thresholding of the signal, watersheding of the binary result before masking with analyzing particles tool. The resulting mask was applied to original image, positive cells quantified manually, and result reported to nuclei automatic quantification for ratio calculation.

### RNA isolation

RNA isolation was performed with RNAqueous-Micro Kit (Invitrogen AM1931) following supplier procedure. For intestinal organoids, cells were retrieved by pipetting with 100 µl of cold lysis buffer per well and transferred to a 1.7 ml conical tube. Lysate was either processed right away following kit procedure or frozen at -20°C until processing. Isolated intestinal epithelial cells were retrieved in 300 µl of lysis buffer by FACS and frozen at -20°C before processing. All samples followed DNase treatment provided by the kit for 30 minutes at 37°C. Total RNA was quantified using a Qubit v3 Fluorometer (Invitrogen Q33216) and the Qubit High Sensitivity RNA Quantification Assay (Invitrogen Q32852). RNA was either frozen at -70°C for genomic downstream analysis or used for RT-qPCR.

### RT-qPCR

20 to 100 ng RNA input was subjected to reverse transcription using the iScript cDNA Synthesis Kit (Bio-Rad 1708891) and diluted 1:5 in molecular grade water (Corning 46-000-CI). RT-qPCR was carried out in technical triplicate using Taqman assays and SsoAdvanced Universal Probes Supermix (BioRad 1725284), following manufacturer protocols and using 1µl cDNA input per reaction at the exception of reactions targeting *Tet1* expression which required 2.5 µl cDNA. Reactions were run on a QuantStudio 3 Real Time PCR instrument (Applied Biosystems A28567) and analyzed using the ddCT method ^52^. *Actb* was selected as the internal housekeeping gene. Taqman assay IDs used in this manuscript are: *Actb* (Mm02619580_g1), *Tet1* (Mm01169087_m1), *ChgA* (Mm00514341_m1), *Muc2* (Mm00458299_m1).

### Omni-ATAC sequencing

Library preparation and sequencing procedure has been adapted from previously described by Corces et al.^38^ and publicly available ATAC-seq protocol from Kaestner lab (2019, Perelman School of Medecine, University of Pennsylvania, Philadelphia, PA, USA). **ATAC reaction.**15 000 *Sox9^low^*intestinal stem cells isolated by FACS were sorted in 500 µl of sorting media. Cells were pelleted at 600g for 5 minutes at 4°C and supernatant immediately and gently discarded. Pellet was resuspended in 50 µl resuspension buffer [10 mM TrisHCl pH 7.4, 10 mM NaCl, 2 mM MgCl_2_] with 0.1% NP40 (ThermoFisher), 0.1% Tween 20 (ThermoFisher) and 0.01% Digitonin (Promega). Suspension was incubated 3 minutes on ice and washed with 1 ml resuspension buffer with 0.1% Tween 20. Nuclei were centrifuged at 500g for 5 minutes at 4°C and supernatant immediately and carefully discarded. Pellet was resuspended in 50 µl of Tn5 transposition mix [25 µl Transposition buffer (Illumina 15027866), 2.5 µl loaded Transposase (Illumina 15027865), 16.5 µl PBS (REF), 0.5 µl 0.1% Digitonin, 0.5 µl 10% Tween 20 and 5 µl mg H_2_O)] and incubated at 37°C and 1000 RPM with a ThermoMixer heating block (Eppendorf). Reaction was stopped with 1:5 volume of gDNA binding buffer from DNA clean & Concentrator 5 kit (Zymo Research D4013). gDNA isolation was performed following manufacturer’s recommendations for fragmented DNA. **Library amplification and indexing.** Pre-amplification and indexing has been carried out in 50 µl reaction mix containing 25 µl NEBNext High-Fidelity 2X PCR Master Mix, 20 µl tagmented gDNA, 2.5 µl 25 µM Ad1-noMX primer and 2.5 µl 25 µM Ad2.x indexing primer with the following PCR program : 72°C 5 minutes, 98°C 30 seconds, 5x [98°C 10 seconds, 63°C 30 seconds, 72°C 1 minute]. The number of cycles to reach saturation has been determined by qPCR in a 20 µl reaction mix containing 5 µl of partially amplified library, 5 µl NEBNext High-Fidelity 2X PCR Master Mix, 3.85 µl mg H2O, 0.5 µl 25 µM Ad1-noMX primer, 0.5 µl 25 µM Ad2.x indexing primer and 0.15 µl 100x SYBR gold with the following program : 98°C 30 seconds, 20x [98°C 10 seconds, 63°C 30 seconds, 72°C 1 minute]. Amplification results were plotted with R against Cycle number in a linear representation. The number of additional PCR cycles needed for each sample has been determined by the number of cycles necessary to reach 1/3 of the maximum R. Amplification was continued with the following program : 98°C 30 seconds, N x [98°C 10 seconds, 63°C 30 seconds, 72°C 1 minute]. Libraries were then cleaned with DNA clean & Concentrator 5 kit (Zymo Research D4013) following manufacturer’s recommendations and eluted in 20 µl mgH_2_O. **Library purification.** Libraries were then purified following double-sided bead purification. Volume was brought up to 40 µl with mgH_2_O and transferred to epi tubes. 0.5X volume of NucleoMag NGS Clean-up and Size Select (Takara Clontech 744970.5) was added and vortexed 5 seconds. After 10 minutes of incubation at RT, epi tubes were placed on magnetic rack for 3 to 4 minutes. Supernatant was transferred to a new tube and 1.3X NucleoMag NGS Clean-up and Size Select of the original library volume was added and tubes vortexed 5 seconds. After 10 minutes of incubation at RT, epi tubes were placed on magnetic rack for 3 to 4 minutes and supernatant discarded. Beads were washed with freshly prepared 200 µl 80% ethanol twice. Beads were span down and tubes left to dry with cap open for 2 to 3 minutes. Beads were resuspended in 15 µl mgH_2_O by pipetting up and down 20 times. Tubes were placed on magnetic rack for 3 to 4 minutes and supernatant transferred to new PCR stripe tubes. **Quality control.** Library concentrations were quantified with Qubit High Sensitivity dsDNA Quantification Assay (Invitrogen Q32851) on a Qubit v3 Fluorometer (Invitrogen Q33216). Library quality and fragment sizes were assessed using High Sensitivity DNA Reagents Kit (Agilent 5067-4626) on a 2100 Bioanalyzer with the support of Emory Integrated Genomics Core. **Sequencing.** Library molarities were determined based on average fragment size and Qubit concentrations. Library pool was set at 3 nM for a final volume of 50 µl and validated by KAPA qPCR. Sequencing was handled by the Emory Integrated Genomics Core on a NextSeq 2000 system (Illumina) with a NextSeq 1000/2000 P2 XLEAP-SBS Reagent Kit (100 Cycles – 2×50 bp paired-end) (Illumina 20100987).

### Single cell RNA-sequencing

For each individual sample 20,000 intestinal epithelial cells were sorted by FACS in RNA lysis buffer for RNA integrity analysis. 500,000 intestinal epithelial cells were sorted for single-cell RNA sequencing in 2 ml of sorting media. Cells were fixed with Evercode Cell Fixation v2 kit (Parse Biosciences ECF2101) following manufacturer’s protocol for whole cells. scRNA-seq libraries were prepared using split-pool barcoding as described in the Evercode WT v2 kit (Parse Biosciences ECW02110). Library was validated by KAPA qPCR and sequenced with the support of the Emory Integrated Genomics Core on a NextSeq 2000 system (Illumina) with a NextSeq 1000/2000 P2 XLEAP-SBS Reagent Kit (200 Cycles – R1: 66; R2: 86; i7:8; i5:8) (Illumina 20100986).

### Genomic data analysis

#### scRNA-seq

The Parse Biosciences bioinformatics pipeline *v.1.1.1* was used to align reads to the reference mouse genome (GRCm39). The output filtered feature counts matrices were further processed and analyzed using Seurat v5.0.1 package ^25^.Low quality cells were filtered out if they contain: less than 500 unique molecular identifiers (UMIs), less than 200 features, or more than 40% mitochondrial UMIs. The Seurat SCTransform method ^53^ was used for normalization and variance stabilization of molecular count data. Clustering was completed by using the RunPCA(), FindNeighbors(), and FindClusters() functions. Nonlinear dimensionality reduction was performed using the uniform manifold approximation and projection (UMAP) technique with the RunUMAP() function implemented in Seurat package. Gene signature enrichment in single-cell data has been realized with UCell package (v3.2.1) and represented either through UMAP projection with FeaturePlot() function or violin plot with VlnPlot() function from Seurat package. Cluster-specific markers and genes differentially expressed between conditions were identified using differential gene expression (DGE) analysis by applying the FindAllMarkers() and FindMarkers() functions with the MAST test ^54^ implemented in the Seurat package. Positive gene set enrichment for cluster characterization has been performed with AddModuleScore () function. Dotplot representation for gene expression across clusters and sub-clusters has been represented with DotPlot() function from package ggplot2 (v3.5.1).

#### Bulk ATAC-seq

Raw ATAC-seq paired-end sequencing reads were processed to remove adapter contamination using *Trimmomatic v.0.39*. Adapter-trimmed reads were then aligned to the mouse genome (GRCm39) using *BWA-MEM v.0.7.19* with default parameters. Duplicate reads were marked with *Picard v.3.1.1 MarkDuplicates* tool. Aligned reads were filtered to retain only properly paired reads with a mapping quality (MAPQ) greater than 5 using *SAMtools v.1.20* was used to remove duplicate reads, as well as the reads with a mapping quality (MAPQ) less than 5 and reads mapped to mitochondrial region. To correct for Tn5 transposase insertion bias, the Binary Alignment Map (BAM) files were processed using *alignmentSieve* tool from *deepTools v.3.5.5* with the *–ATACshift* option. This step offsets reads by +4 bp on the positive strand and −5 bp on the negative strand to reflect the actual cut sites. The shifted BAM files were used to generate normalized coverage tracks (bigWig files) using *deepTools bamCoverage*, with normalization set to reads per genomic content (RPGC). Peaks representing accessible chromatin regions were identified using *MACS2 v.2.2.9.1* in narrow peak mode. The differential binding analysis was done using *DiffBind v 3.16.0* package and peaks were annotated using *ChIPseeker v.1.42.1* package.

### Statistical analysis

Statistical analyses were carried out in Prism 10.4.2 (GraphPad software). For statistical comparison, unpaired t-test was used for single feature comparison between control and Tet1iKO cohorts. Ordinary one-way ANOVA was conducted to compare a single feature in several conditions between control and Tet1iKO. A p value <0.05 was set as threshold for significance. All values are depicted as mean ± SEM unless indicated differently. Significance between two sets of measures is indicated on representations with * for p-value <0.05, ** p < 0.01, *** p < 0.001. Significance between more than 2 sets of measures is indicated alphabetically.

## Author contributions

ARG, NVJ, and ADG conceived and designed the experiments. ARG, NVJ, VR, GA, SS, MF and ADG performed experiments. ARG, NVJ, VR, and SS analyzed data. ARG and SB carried out computational analyses. ARG interpreted results of experiments, prepared figures, and drafted manuscript. ARG and ADG edited and revised manuscript. ADG approved final version of manuscript.

## Acknowledgements

We thank Dr. Scott Magness and members of the Magness lab (UNC) for providing *Sox9^EGFP^* and *VilCreER* mice. We thank Dr. Ajana Rao and members of the Rao lab (LJI) for providing *Tet1^fl/fl^* mice. We thank Dr. David Gorkin and members of the Gorkin lab (Emory) for helpful discussions and feedback as well as sharing reagents for experimental support. We thank Dr. Bing Yao and members of the Yao lab (Emory) for helpful discussions and feedback as well as providing reagents for experimental support. We thank the members of the Gracz lab for constructive discussion and critical reading of the manuscript. Research reported in this publication was supported in part by the Emory University Integrated Genomics Core (EIGC; RRID:SCR_023529) of the Winship Cancer Institute of Emory University and NIH/NCI under award number, 2P30CA138292-04. This work was supported by the Emory University Integrated Cellular Imaging Core Facility (ICI; RRID:SCR_023534).

## Notes

This study was funded by the NIH/NIGMS under award numbers R35GM142503 (Gracz) and T32GM008490 (Boss), and by the NIH/NIDDK under award number F31DK136254 (Janto). Research reported in this publication was supported in part by the Emory University Emory Integrated Cellular Imaging Core Facility (RRID:SCR_023534) and NIH under Award Number S10 OD032320-01. The content is solely the responsibility of the authors and does not necessarily represent the official views of the National Institutes of Health.

### Competing Interest Statement

The authors have declared no competing interest.

