## Supplementary figures for "Tet1 safeguards lineage allocation in intestinal stem cells"

Figure S1 – Tet1iKO confirmation from isolated intestinal stem cells.

Figure S2 – Tet1iKO phenotypes are variable across housing conditions.

Figure S3 – scRNA-seq clustering parameters for conventional housing.

Figure S4 – scRNA-seq clustering parameters for high-level barrier housing.

Figure S5 – Global transcriptomic changes in Tet1iKO IECs are more pronounced in CONV housing.

Figure S6 – Differential expression analysis between control and Tet1iKO mice in CONV and HL housing.

Figure S7 – scRNA-seq clustering parameters for ISC subclustering.

Figure S8 – Gating strategy for intestinal epithelial cell sorting.

Figure S9 – Gating strategy for Sox9$^{EGFP}$ populations.

**Supplementary data**

Table S1 – IEC gene sets used for annotating scRNA-seq.

Table S2 – Differential gene expression analysis results for CONV housing.

Table S3 – Differential gene expression analysis results for HL housing.

Table S4 – Differential peak analysis (DiffBind) results for ATAC-seq.

Table S5 – TOBIAS results for ATAC-seq.

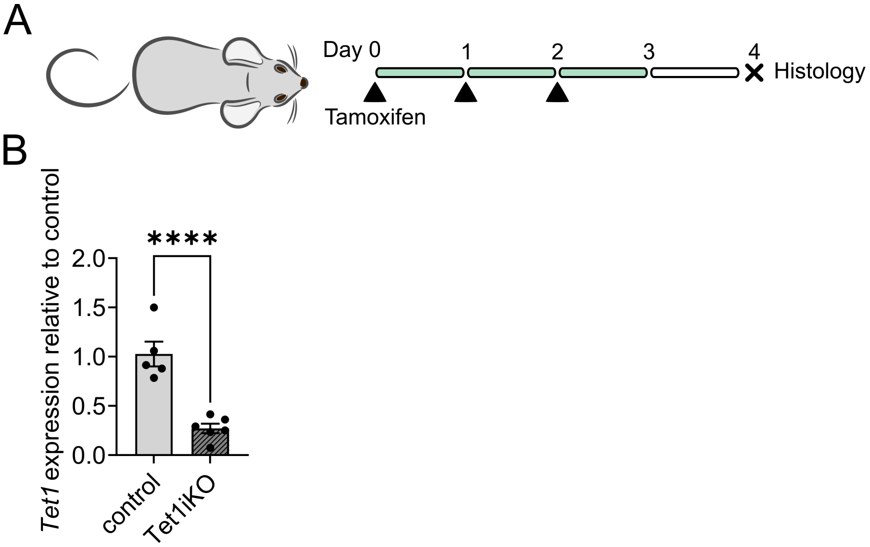


***Figure S1 – Tet1iKO confirmation from isolated intestinal stem cells.***

(A) *Tet1* recombination (Tet1iKO) is induced by daily tamoxifen injection for 3 days followed by 2 days washout. Control animals harbor the *VilCreER* transgene and are wild-type for *Tet1*. (B) RT-qPCR demonstrates a significant reduction of *Tet1* expression in Sox9$^{low}$ ISCs isolated from Tet1iKO vs control mice (n = 5 control and 6 Tet1iKO mice. **** indicates significance, p < 0.0001).

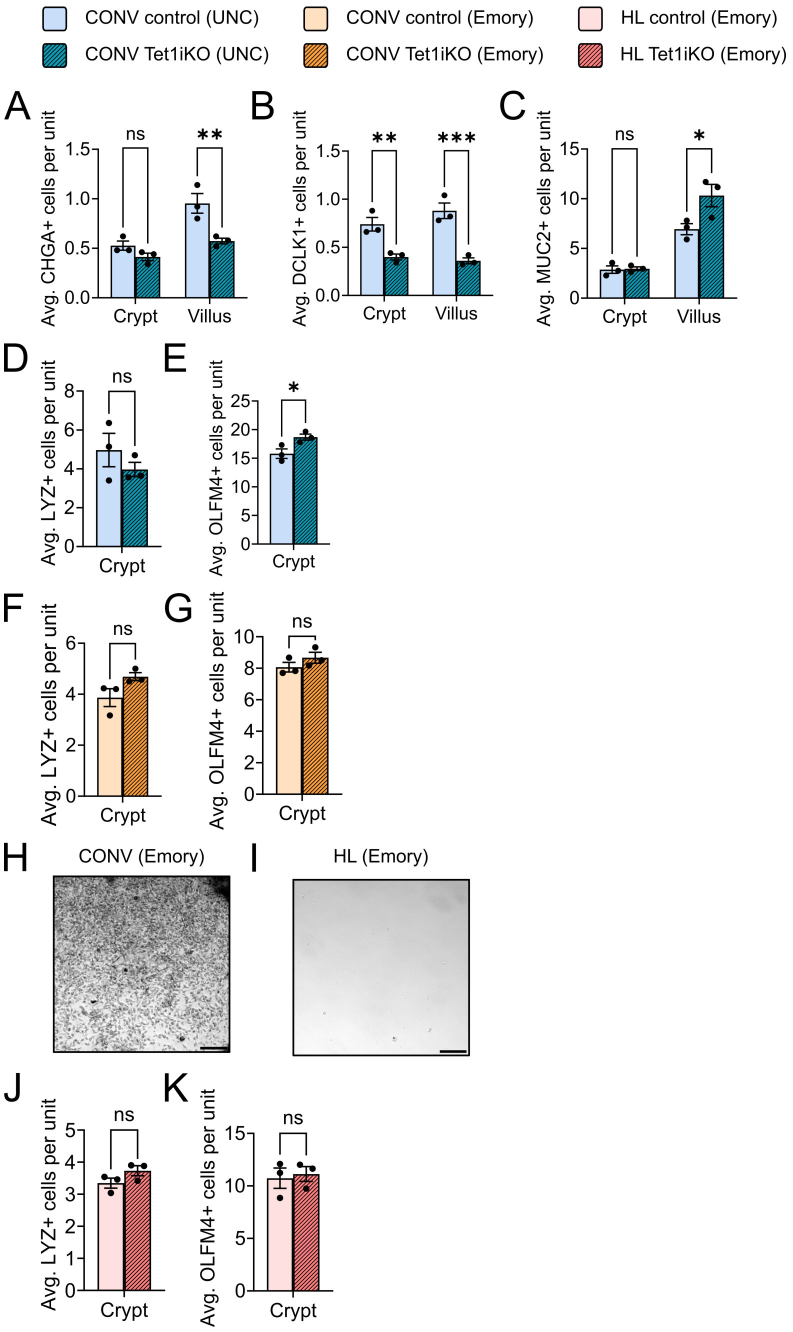


***Figure S2 – Tet1iKO phenotypes are variable across housing conditions.***

IF quantification in CONV (UNC) housing revealed (A) a significant decrease in enteroendocrine cell (CHGA) number in Tet1iKO villi, (B) a significant decrease in tuft cell (DCLK1) number in Tet1iKO crypts and villi, (C) a significant increase in goblet cell (MUC2) number in Tet1iKO villi, and (D) no change in Paneth cell (LYZ) number. (E) Numbers of crypt-based cells expressing ISC marker OLFM4 were significantly increased in Tet1iKO from CONV (UNC) housing. In CONV (Emory) housing, (F) Paneth cell numbers remained unchanged between Tet1iKO and controls and (G) no differences were observed in numbers of OLFM4+ cells (H) Brightfield microscopy of mouse cecal contents reveals the presence of commensal protist *Tmu* in CONV (Emory) housing (scale bar represents 100 µm). (I) Cecal contents from mice housed in HL (Emory) are negative for *Tmu* (scale bar 100 µm). (J) Similar to CONV (UNC) and CONV (Emory) samples, mice in HL (Emory) housing exhibit no difference in Paneth cell numbers. (K) OLFM4+ cell numbers are also unchanged between HL (Emory) Tet1iKO and control intestines (control n = 3, Tet1iKO n = 3, * indicates $p<0.05$, ** for $p<0.01$ and *** for $p<0.001$).

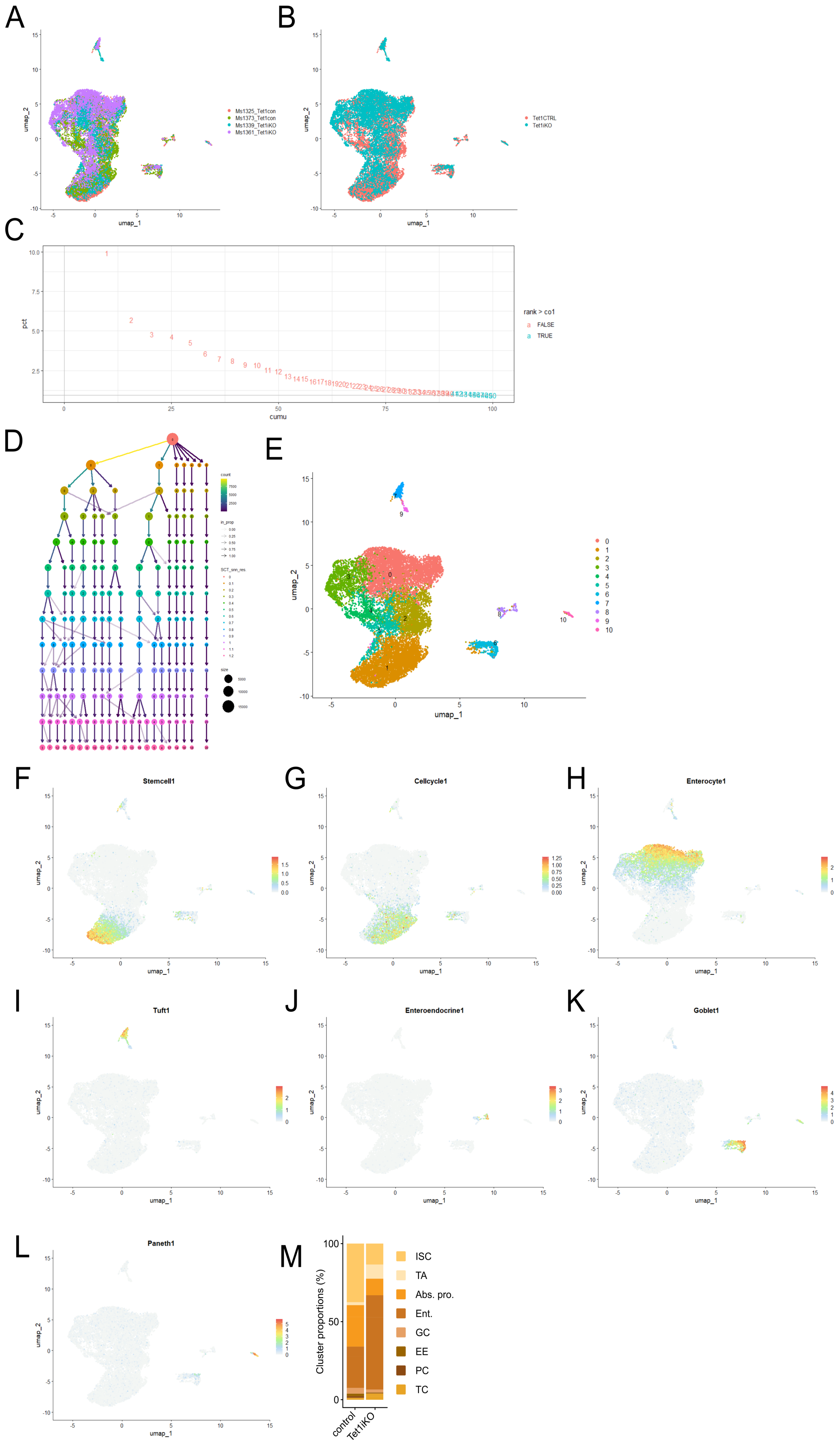
A
B
C
D
E
F
G
H
I
J
K
L
M
umap_1
umap_2
Ms1325_Tet1con
Ms1373_Tet1con
Ms1339_Tet1iKO
Ms1361_Tet1iKO
Tet1CTRL
Tet1iKO
pct
cumu
rank > co1
FALSE
TRUE
count
in_prop
SCT_snn_res
size
Stemcell1
Cellcycle1
Enterocyte1
Tuft1
Enteroendocrine1
Goblet1
Paneth1
Cluster proportions (%)
control
Tet1iKO
ISC
TA
Abs. pro.
Ent.
GC
EE
PC
TC

***Figure S3 – scRNA-seq clustering parameters for conventional housing.***

(A) UMAP of individual sample distribution from scRNA-seq in CONV housing. (control n = 2, Tet1iKO n = 2) (B) UMAP of control and Tet1iKO samples from scRNA-seq analysis in CONV housing demonstrates equal representation and distribution of groups. (C) 40 PCs were retained under the cut-off of 90% of the cumulative variance in CONV housing scRNA-seq analysis (cumulative variance, red < 0.9 and green > 0.9). (D) 11 clusters were identified with a resolution of 0.3 based on clustering analysis in Clustree. (E) UMAP representing 11 identified clusters with 40 PC and a resolution of 0.3. (F-L) UMAPs representing distribution of IEC subpopulation defining gene expression, used to annotate cluster identity (list of genes used is provided in Table S1). (M) Cluster proportions demonstrate a greater proportion of Ent. population in Tet1iKO samples, along with decreased relative numbers of ISC, TA, and Abs. pro.

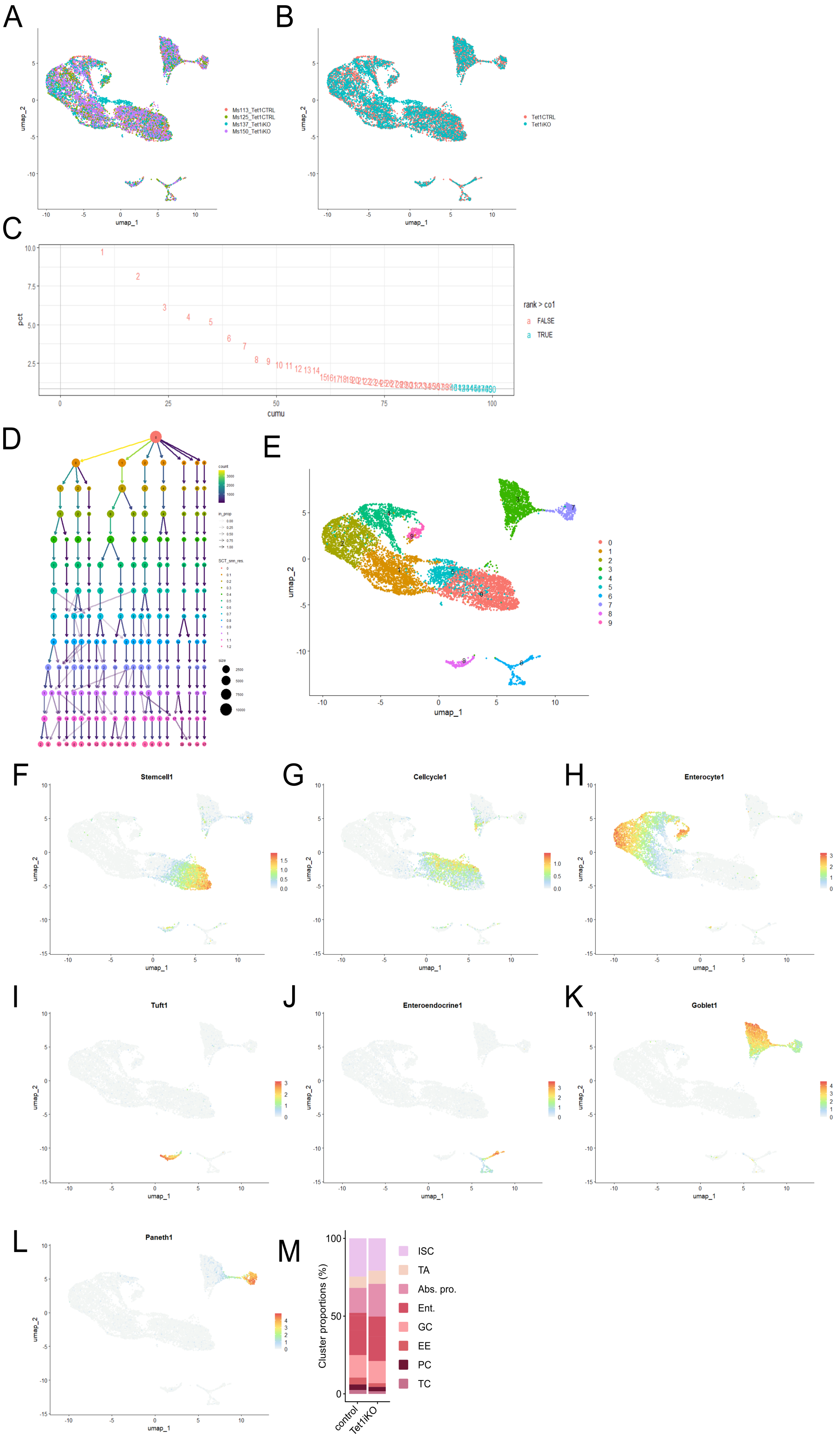
A
Ms113_Tet1CTRL
Ms125_Tet1CTRL
Ms137_Tet1iKO
Ms150_Tet1iKO
umap_1
umap_2
B
Tet1CTRL
Tet1iKO
C
pct
cumu
rank > co1
FALSE
TRUE
D
count
in_prop
SCT_snn_res
size
E
F
Stemcell1
G
Cellcycle1
H
Enterocyte1
I
Tuft1
J
Enteroendocrine1
K
Goblet1
L
Paneth1
M
Cluster proportions (%)
control
Tet1iKO
ISC
TA
Abs. pro.
Ent.
GC
EE
PC
TC

***Figure S4 – scRNA-seq clustering parameters for high-level barrier housing.***

(A) UMAP of individual sample distribution from scRNA-seq analysis across HL housing. (control n = 2, Tet1iKO n = 2) (B) UMAP of control and Tet1iKO samples from scRNA-seq analysis in HL housing demonstrates equal representation and distribution of groups. (C) 39 PCs were retained under the cut-off of 90% of the cumulative variance in CONV housing scRNA-seq analysis (cumulative variance, red < 0.9 and green > 0.9). (D) 10 clusters were identified with a resolution of 0.3 based on clustering analysis in Clustree. (E) UMAP representing 10 identified clusters with 39 PC and a resolution of 0.3. (F-L) UMAPs representing distribution of IEC subpopulation defining gene expression, used to annotate cluster identity (list of genes used is provided in Table S1). (M) Cluster proportions demonstrate few changes in IEC populations between Tet1iKO and control.

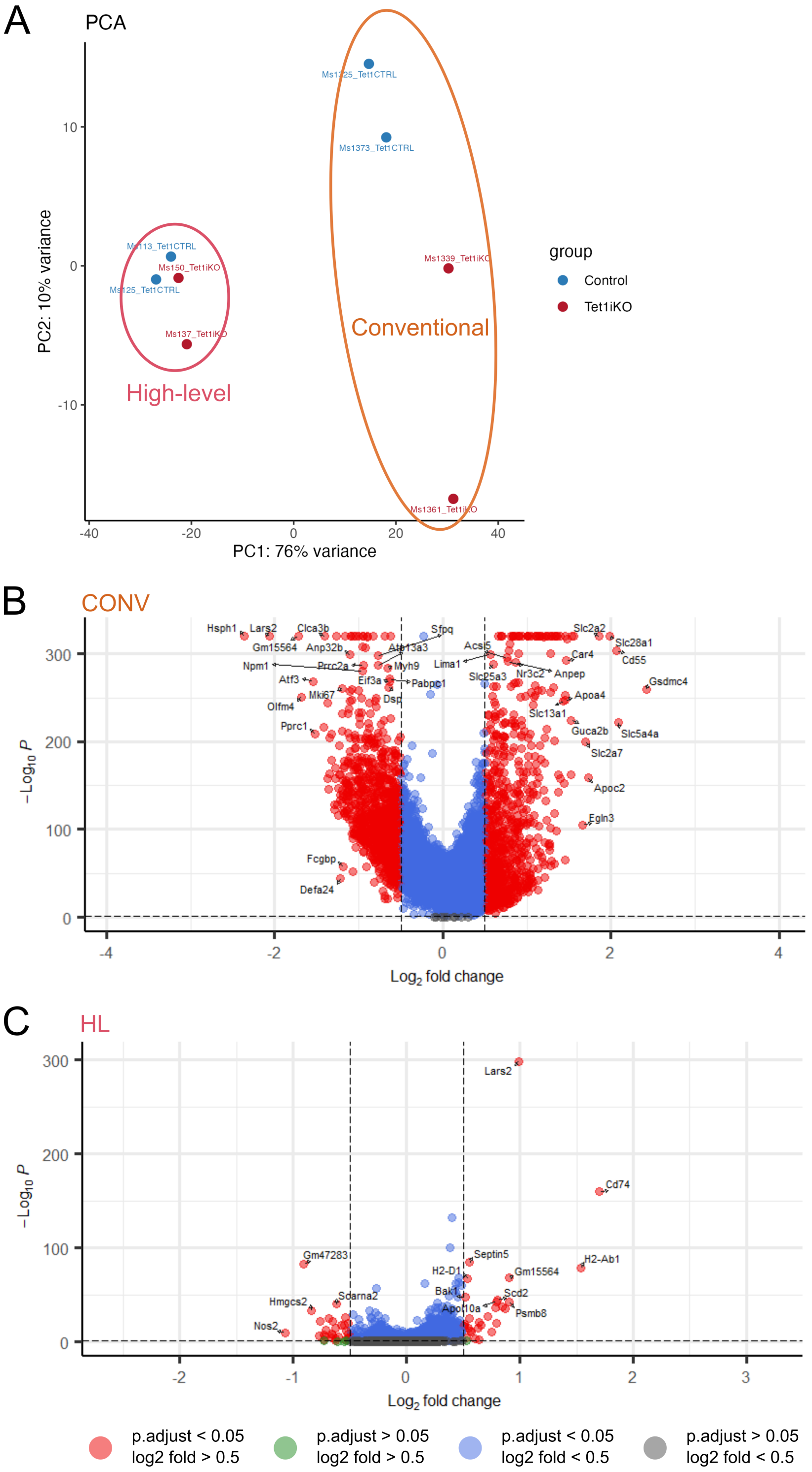


***Figure S5 – Global transcriptomic changes in Tet1iKO IECs are more pronounced in CONV housing.***

(A) PCA demonstrates that IEC transcriptomes from HL and CONV housing primarily cluster based on their housing condition, but that greater differences between control and Tet1iKO groups are present in CONV housed mice. (B) Differential expression analysis using MAST identifies a significant number of DEGs from scRNA-seq on CONV housed Tet1iKO IECs across all subpopulations. (C) MAST identifies significantly fewer DEGs across all IECs in HL housed Tet1iKO mice, consistent with decreased transcriptomic differences identified by PCA (grey dots = log2 fold change < 0.5, p-adjusted value <0.05 ; blue dots = log2 fold change > 0.5, p-adjusted value < 0.05 ; red dots = log2 fold change > 0.5, p-adjusted value > 0.5).

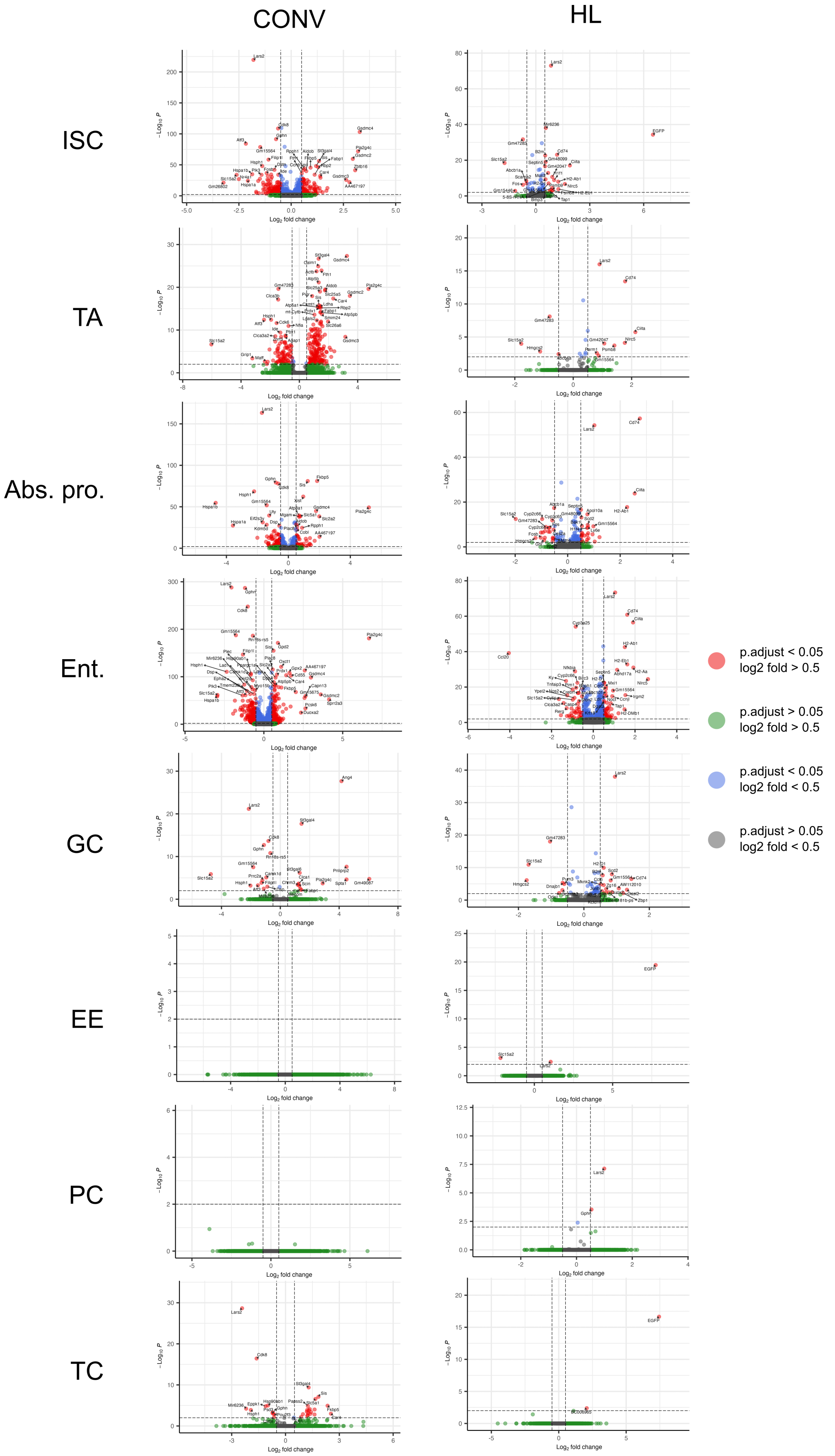
CONV
HL
ISC
TA
Abs. pro.
Ent.
GC
EE
PC
TC
p.adjust < 0.05
log2 fold > 0.5
p.adjust > 0.05
log2 fold > 0.5
p.adjust < 0.05
log2 fold < 0.5
p.adjust > 0.05
log2 fold < 0.5

***Figure S6 – Differential expression analysis between control and Tet1iKO mice in CONV and HL housing.***

Differential expression analysis between control and Tet1iKO mice reveals that the number of DEGs are overall more numerous in scRNA-seq data from CONV compared to HL housing. “ISC”, “TA”, “Abs.pro.” and “Ent.” clusters demonstrate the most DEGs in both housing conditions (CONV and HL) (Table S2 and S3).

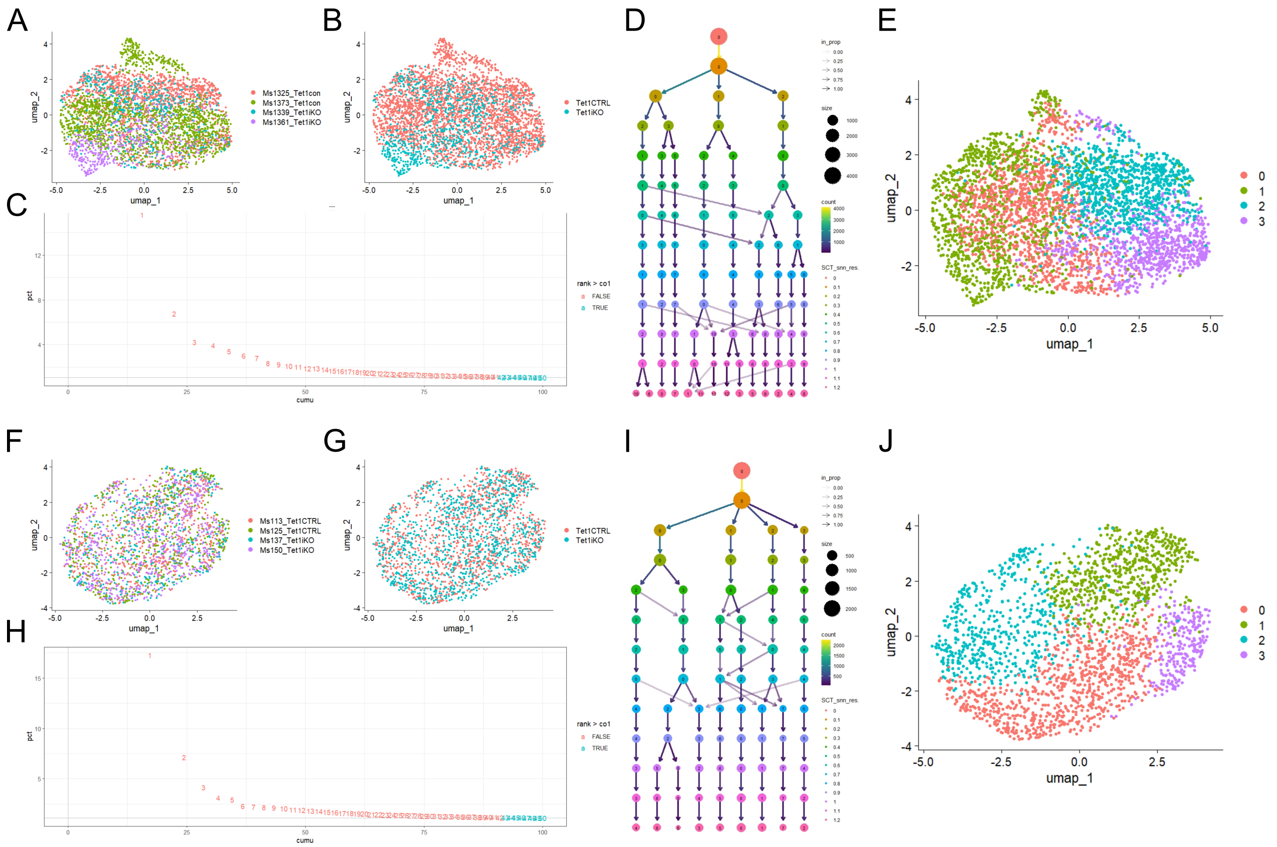


***Figure S7 – scRNA-seq clustering parameters for ISC subclustering.***

(A) UMAP of individual sample distribution from scRNA-seq analysis across CONV housing (control n = 2, Tet1iKO n = 2). (B) UMAP of ISCs from CONV scRNA-seq demonstrates differential distribution of Tet1iKO and control groups. (C) 41 PCs were retained under the cut-off of 90% of the cumulative variance in sub-analyzed ISCs from CONV scRNA-seq (cumulative variance, red < 0.9 and green > 0.9). (D) 4 clusters were identified with a resolution of 0.3 based on clustering analysis in Clustree. (E) UMAP representing 4 identified ISC subclusters with 41 PC and a resolution of 0.3. (F) UMAP of individual sample distribution from scRNA-seq analysis across HL housing. (control n = 2, Tet1iKO n = 2) (F) UMAP of ISCs from HL scRNA-seq demonstrates equal distribution of Tet1iKO and control groups. (H) 42 PCs were retained under the cut-off of 90% of the cumulative variance in sub-analyzed ISCs from HL scRNA-seq (cumulative variance, red < 0.9 and green > 0.9). (D) 4 clusters were identified with a resolution of 0.3 based on clustering analysis in Clustree. (E) UMAP representing 4 identified ISC subclusters with 42 PC and a resolution of 0.3.

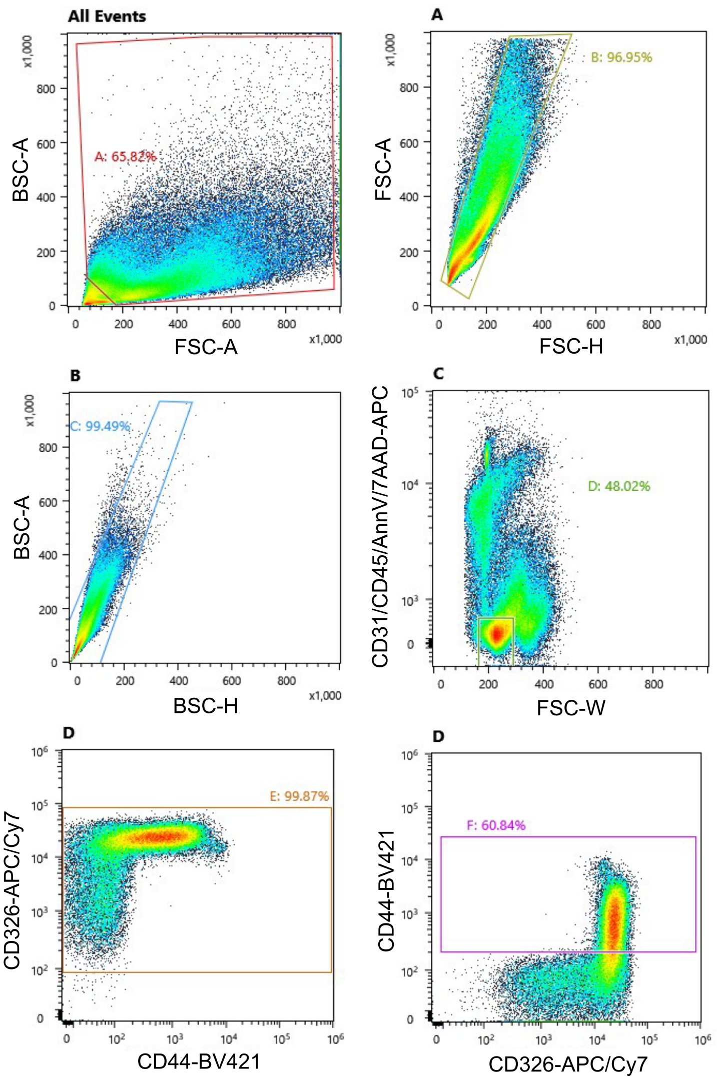


***Figure S8 – Gating strategy for intestinal epithelial cell sorting.***

FACS gating strategy to isolate total IECs for scRNA-seq experiments.

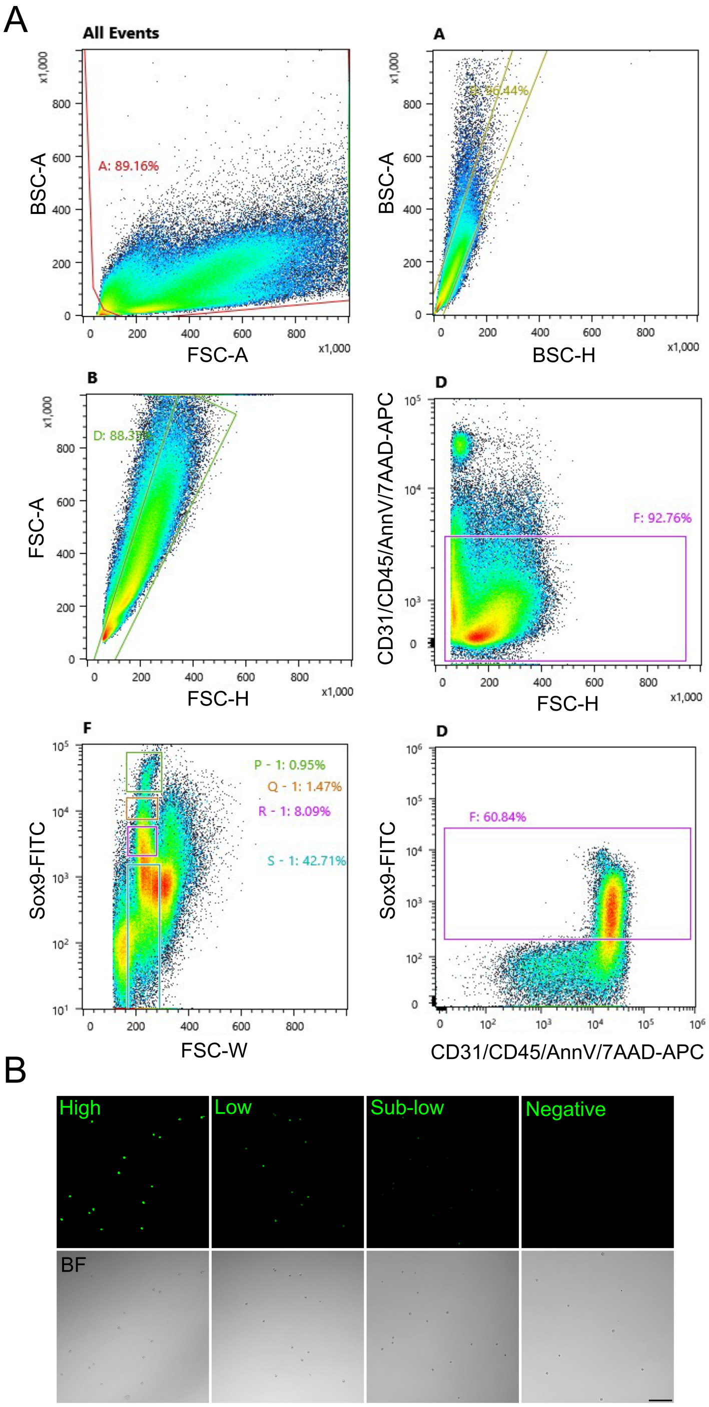


***Figure S9 – Gating strategy for Sox9$^{EGFP}$ populations.***

(A) FACS gating strategy to isolate Sox9$^{low}$ ISCs for single cell organoid assays and ATAC-seq. (B) Representative pictures of isolated Sox9$^{EGFP}$ populations show distinct EGFP levels from High to negative EGFP cells (scale bar represents 50 μm).
